# Experimentally Defined Tm-Guided Exon–Exon Junction Primer Design Enables Specific Discrimination of Highly Similar RNA Variants

**DOI:** 10.64898/2026.04.02.716213

**Authors:** Junyeop Ahn, Yun H. Lee, Donald J Zack, Ping-Wu Zhang

**Author notes:** These authors contributed equally to this work (Junyeop Ahn, Yun H. Lee). **Corresponding author:** Ping-Wu Zhang Contact Address: Wilmer Eye Institute, Johns Hopkins University School of Medicine, 400 N Broadway, Smith Building, Room 3001, Baltimore, Maryland, 21231, USA.

## Abstract

Accurate detection of individual RNA transcript variants is challenging when closely related transcripts share extensive exonic sequence and differ primarily in exon connectivity. Analysis of curated human RefSeq transcripts identified 9,546 protein-coding genes containing at least one high-similarity RNA variant, highlighting the need for transcript-discriminating assays beyond conventional exon-targeted PCR. We developed a melting-temperature (Tm)-guided exon-exon junction (EEJ) RT-PCR strategy in which the 5’ and 3’ exon-matching arms of a junction primer are evaluated independently. Systematic experiments showed that productive amplification was lost when a partially matched primer segment had a Tm approximately 13-16 °C below the PCR annealing temperature, supporting an approximately 15 °C arm-specific design criterion. Tm-guided EEJ primers, including designs generated with the automated TransPrimer tool, specifically amplified highly similar transcript variants of HTRA1-AS1, APEX1, and GAPDH, reduced heteroduplex formation, and produced single qPCR melting peaks; most primer pairs showed 90-110% efficiency with R^2^ > 0.98. This approach provides an experimentally grounded framework for transcript-specific RT-PCR and qPCR when distinctive exonic regions are limited.

## Introduction

Alternative splicing is a major source of transcriptomic and proteomic diversity in eukaryotic organisms. Human genes commonly contain multiple exons and generate multiple transcript isoforms through mechanisms including alternative exon usage, exon skipping, alternative splice-site selection, and intron retention. These transcript variants can differ in regulatory function, subcellular localization, and protein-coding potential and can contribute to development, disease progression, and cellular responses to stress [1–4]. Genome-wide analyses have demonstrated extensive transcript isoform diversity across human genes, and increasingly comprehensive transcript annotations and long-read sequencing continue to reveal substantial transcript complexity [5–8]. However, transcript multiplicity alone does not capture the sequence and structural relationships among sibling isoforms. In particular, transcript variants may share extensive exonic sequence and differ primarily in exon connectivity, creating highly similar transcript structures that can be difficult to distinguish experimentally.

Reliable detection and quantification of individual transcript variants therefore remain technically challenging, particularly when related transcripts share extensive exonic sequence [9, 10]. Conventional RT-PCR and quantitative PCR assays typically rely on primers targeting transcript-specific exonic regions or on differences in amplicon size. However, many transcript variants lack sufficiently large unique exonic regions and instead differ mainly in exon connectivity. Under these conditions, exon-targeted primer designs can coamplify multiple transcripts and compromise variant-specific detection [9, 10]. Despite extensive characterization of transcript multiplicity across the human transcriptome [5–8], the prevalence of structurally high-similarity variants and the resulting challenge for transcript-specific PCR assay design have not been systematically quantified.

High-throughput sequencing approaches, including short-read RNA sequencing and long-read sequencing, have greatly improved transcriptome-wide isoform resolution [11, 12]. Nevertheless, these methods can be costly, computationally intensive, and impractical for routine validation or targeted quantification of a limited number of transcripts. PCR-based assays therefore remain widely used for validating splice events and measuring transcript abundance in experimental studies [13,14].

A complementary strategy is to exploit exon–exon junctions created by RNA splicing. Because transcript variants are defined in part by their specific combinations of exon junctions, primers that span a junction can provide transcript specificity even when large unique exonic regions are absent. Exon-junction and intron-spanning primer designs are also widely used to reduce amplification from genomic DNA [13,14]. Several computational resources support junction-primer or isoform-aware primer design, including ExonSurfer, Ex-Ex Primer and Primer-BLAST [15–17]. These resources provide important capabilities for transcript selection, junction placement, specificity assessment, or primer optimization. However, junction design is commonly expressed through nucleotide-overlap, primer-length, GC-content, and overall-Tm constraints, while a systematically established experimental relationship between the Tm of each partially matched junction-primer arm and productive amplification has not been defined.

Here, we describe an experimentally defined Tm-guided EEJ RT-PCR strategy and utilization of an accompanying web tool, TransPrimer, for discriminating closely related RNA transcript variants [18]. The method treats the two exon-matching segments of a junction-spanning primer as independent thermodynamic recognition arms. Each arm is designed to remain insufficiently stable on its own under the selected annealing conditions, whereas the full-length primer remains stable when both arms anneal across the correct exon junction. We systematically establish an arm-specific Tm criterion, evaluate the method using RT-PCR, qPCR, and Sanger sequencing, and examine its effect on heteroduplex formation.

## Materials and Methods

### Human transcript variant analysis and identification of structurally non-unique transcripts

Human protein-coding genes and their associated transcript annotations were retrieved from the National Center for Biotechnology Information (NCBI) RefSeq database [19]. A total of 20,593 human protein-coding genes were included in the initial dataset. Transcript records associated with each gene were collected from RefSeq and classified according to accession type as curated protein-coding mRNA transcripts (NM), curated non-coding RNA transcripts (NR), computationally predicted mRNA transcripts (XM), and computationally predicted non-coding RNA transcripts (XR). Of the 20,593 input genes, 20,583 were successfully processed. The composition of the resulting transcript dataset and the distribution of curated NM transcripts among genes are summarized in Fig. 1A and Fig. 1B, respectively.

**Figure 1.**
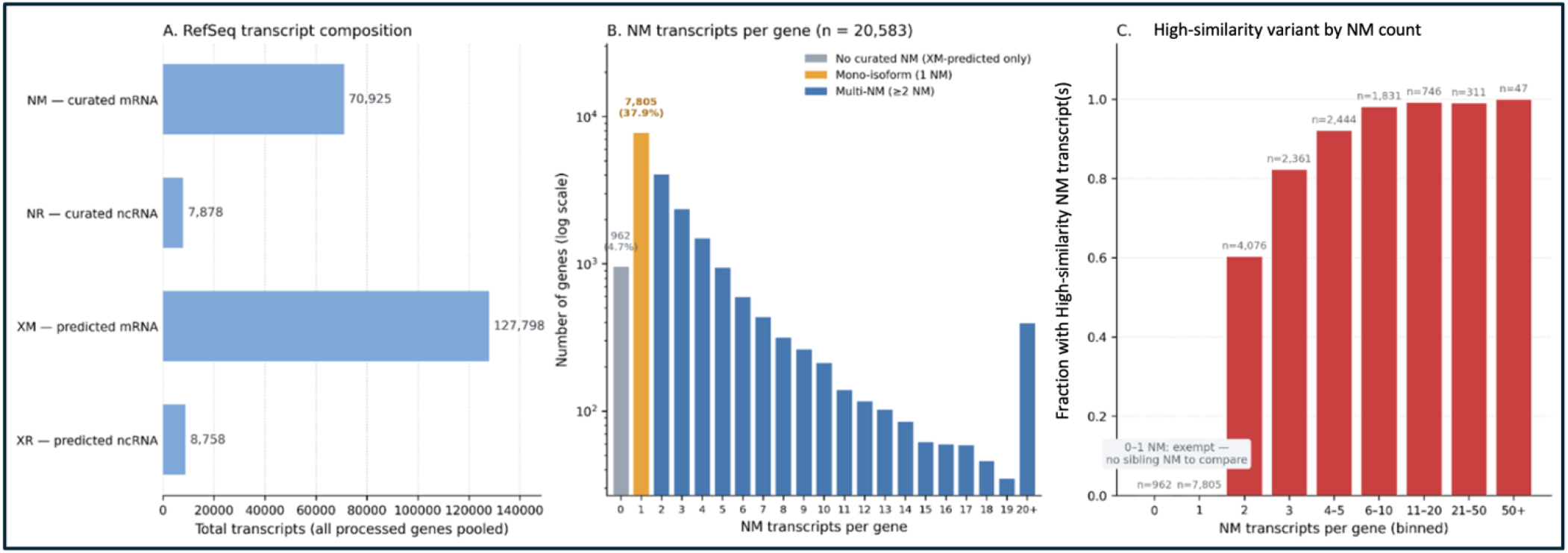
Curated and predicted transcript landscape of human protein-coding genes. **(A)** RefSeq transcript composition across 20,583 successfully processed human protein-coding genes (GRCh38), including 70,925 curated NM, 7,878 curated NR, 127,798 predicted XM, and 8,758 predicted XR transcripts. (B) Distribution of genes by curated NM transcript number. Of the processed genes, 962 (4.7%) lacked curated NM transcripts, 7,805 (37.9%) contained a single NM transcript, and the remainder contained ≥2 NM transcripts. (C) Prevalence of high-similarity RNA variants according to NM transcript number. A transcript was classified as high-similarity when every exon was fully contained within an exon of the same sibling NM transcript. Genes with zero or one NM transcript were exempt from comparison. The prevalence of high-similarity variants increased with transcript number, approaching 100% in transcript-rich genes.

Structural transcript comparisons were restricted to curated RefSeq (NM) transcripts mapped to the GRCh38 reference assembly. Genes lacking a curated NM transcript were excluded, and single-NM genes were considered non-comparable because of the absence of sibling transcripts for within-gene discrimination. Genes harboring two or more curated NM transcripts were classified as multi-NM genes and subjected to pairwise exon-structure comparisons.

Within each multi-NM gene, every curated transcript was evaluated pairwise and directionally against each sibling transcript. A transcript was operationally defined as a “high-similarity variant” if every exon within it was fully contained within exons of a sibling transcript. A gene was classified as containing a high-similarity variant if at least one of its NM transcripts met this complete exon-containment threshold.

We evaluated the frequency of genes containing high-similarity variants across strata defined by the number of curated NM transcripts per gene. Under this framework, the high-similarity designation denotes a transcript configuration lacking any structurally exclusive exons relative to at least one sibling isoform. Consequently, selective amplification of such variants cannot rely on unique exonic sequences, requiring alternative discrimination strategies such as exon–exon junction-targeted primer placement.

### Primer design

For the HTRA1-AS1 experiments used to establish and characterize the arm-specific Tm rule, primer candidates were initially generated using Primer3 software [20], and EEJ primers were refined manually. After the approximately 15 °C arm-specific criterion was established, the design logic was implemented in TransPrimer [18]. Transcript-specific primer pairs for the APEX1 and GAPDH qPCR experiments were generated using the automated TransPrimer workflow, which compares the selected transcript with sibling RefSeq variants, identifies discriminating exon or EEJ features, evaluates single- and dual-EEJ strategies when required, and generates compatible partner primers. For each junction-spanning candidate, the complete primer and its two exon-matching arms are evaluated independently so that the experimentally derived arm-specific Tm constraint can be applied during automated design. Detailed software implementation and computational evaluation are described in the accompanying TransPrimer manuscript [18].

Primer lengths were typically 18–24 nucleotides, with melting temperatures (Tm) of approximately 58–64 °C and GC content between 40–60%. When sequence constraints limited GC content, primer length was adjusted to maintain appropriate Tm. Amplicon sizes were designed to range from 150 to 250 bp to ensure efficient amplification and compatibility with both conventional RT-PCR and quantitative PCR. Primer specificity was evaluated against reference genomic and transcript sequences using BLAST to minimize off-target amplification. When multiple transcript variants shared similar junctions, additional primers targeting alternative junctions were designed to improve discrimination. All primers were synthesized by IDT (Integrated 156 DNA Technologies, Coralville, IA, USA).

### PCR amplification and optimization

For HTRA1-AS1 primer optimization and the empirical Tm-series experiments, primer melting temperatures (Tm) were calculated using the NEB Tm Calculator (v1.16.10), with Phusion High-Fidelity DNA Polymerase under HF buffer conditions as the reference. For each junction-spanning primer, the Tm values of the 5′- and 3′-exon-matching arms were calculated separately, together with the full-length primer Tm, to guide primer design and PCR optimization. For the APEX1 and GAPDH primers generated by TransPrimer, the software independently calculates the full-length and arm-specific Tm values using its implemented thermodynamic model and user-defined reaction conditions, as described in the TransPrimer manuscript [18].

PCR amplification was performed using Phusion High-Fidelity PCR Master Mix (Thermo Fisher Scientific, USA) or PyroMark PCR Master Mix (Qiagen, USA) according to the manufacturers’ instructions. PCR conditions were optimized to ensure that the partial Tm values of junction-spanning primers remained below the annealing temperature, thereby minimizing nonspecific amplification.

### Sanger sequencing validation

PCR products corresponding to each transcript variant were validated by Sanger sequencing at the Johns Hopkins Genetic Resources Core Facility. PCR products were submitted without purification using 2 µL PCR product, 1 µL of 10 µM sequencing primer, and 7 µL H2O. Sequencing chromatograms were analyzed and aligned to reference sequences to confirm the identity of the amplified products. This validation step verified that each amplicon corresponded precisely to the intended exon–exon junction, confirming the specificity and reliability of the primer design and amplification strategy.

### Determination of optimal arm-specific Tm constraints for EEJ primers

A first primer series sharing HTRA1-AS1 target-complementary sequence but containing progressively shortened target-matching regions was tested with a common partner primer to determine the relationship between primer Tm and amplification at a fixed annealing temperature. A second series retained the same HTRA1-AS1-complementary 3′ arm but replaced the 5′ arm with random noncomplementary sequences and was tested across annealing temperatures of 58, 60, 62, and 64 °C. These experiments were used to determine whether a partially matched arm could independently support productive extension and to establish a practical Tm difference between the annealing temperature (Ta) and the Tm of an individual exon-matching arm. Reactions were performed using iTaq™ Universal SYBR® Green Supermix (Bio-Rad, USA, Cat. No. 1725120) for 40 cycles.

### Quantitative PCR and Assessment of Primer Efficiency

RNA was extracted from HEK293T cells using the RNeasy® Mini Kit (Qiagen, USA, Cat. No. 74104) according to the manufacturer’s instructions. cDNA was synthesized from 1 µg of total RNA using the iScript™ cDNA Synthesis Kit (Bio-Rad, USA, Cat. No. 1708890). HTRA1-AS1 primer pairs were based on the experimentally optimized EEJ designs described above, whereas transcript-specific APEX1 and GAPDH primer pairs were generated using TransPrimer, incorporating the experimentally derived arm-specific Tm constraint of approximately 15 °C below the annealing temperature.

For qPCR melting-curve analysis, reactions containing the designed primer pairs were prepared using cDNA corresponding to 8.3 ng of input RNA and performed in 384-well plates. Each 5-µL reaction contained 2.5 µL of iTaq™ Universal SYBR® Green Supermix and 0.5 µL of 5 µM primer mix. To evaluate primer efficiency, a five-point, 10-fold serial dilution of cDNA was prepared, with the highest concentration corresponding to 16.6 ng of input RNA equivalent per reaction. All qPCR reactions were performed using iTaq™ Universal SYBR® Green Supermix (Bio-Rad, Cat. No. 1725120) on a CFX384 Touch Real-Time PCR Detection System (Bio-Rad). Amplification and melting curves were analyzed using CFX Maestro Software, and amplification efficiencies were calculated from standard curves generated using the serially diluted cDNA samples. Experimental reporting and interpretation of amplification specificity, efficiency, and linearity were guided by established qPCR quality principles, including the MIQE framework [21].

## Results

### Coordinate-based transcript characterization reveals widespread occurrence of high-similarity RNA variants

Analysis of 20,583 successfully processed human protein-coding genes revealed substantial variation in the number and type of RefSeq transcripts associated with individual genes. In total, the dataset contained 70,925 curated NM mRNA transcripts, 7,878 curated NR non-coding RNA transcripts, 127,798 predicted XM mRNA transcripts, and 8,758 predicted XR non-coding RNA transcripts (Figure1A). Predicted XM transcripts were approximately 1.8-fold more abundant than curated NM transcripts, indicating that the numerical composition of the RefSeq transcriptome differs substantially depending on whether curated or computationally predicted records are considered. To maintain a well-defined and consistently curated transcript set, subsequent structural analyses were therefore restricted to curated NM transcripts.

The number of curated NM transcripts varied markedly among protein-coding genes (Figure 1B). Of the 20,583 successfully processed genes, 962 (4.7%) contained no curated NM transcript, although predicted XM transcripts were present, and were therefore excluded from within-gene NM transcript comparisons. An additional 7,805 genes (37.9%) contained only a single curated NM transcript and thus lacked a sibling NM transcript for structural comparison. The remaining 11,816 genes (57.4%) contained two or more curated NM transcripts and constituted the population subjected to pairwise exon-containment analysis.

The prevalence of high-similarity RNA variants increased markedly with transcript multiplicity (Figure 1C). Among genes containing exactly two curated NM transcripts, approximately 60% contained at least one high-similarity RNA variant. This proportion increased to approximately 82% among genes with three NM transcripts and approximately 92% among genes with four to five NM transcripts. The prevalence increased further among transcript-rich genes, reaching approximately 98% for genes containing 6–10 curated NM transcripts and approaching 100% among genes with larger numbers of NM variants. Thus, the probability that a gene contains at least one high-similarity RNA variant increased strongly with the number of curated NM transcripts, demonstrating that structurally similar transcript variants are particularly prevalent among genes with extensive curated transcript diversity.

The structural criterion used to define high-similarity RNA variants is illustrated at exon-level resolution for GAPDH (Figure 2). Five curated NM transcripts were identified for this gene. Three transcripts— NM_001289745.3, NM_001357943.2, and NM_002046.7—were classified as high-similarity RNA variants because every exon of each transcript was fully contained within an exon of the same sibling transcript, NM_001289746.2. In contrast, NM_001256799.3 and NM_001289746.2 did not satisfy the high-similarity criterion. NM_001256799.3 contains a distinct 5′ exon configuration that is not fully represented within a single sibling transcript, whereas the extended 5′ exon of NM_001289746.2 is not fully contained within the corresponding exon of the same sibling transcript. This example illustrates the directional nature of the classification: a transcript is designated as a high-similarity variant only when all of its exons satisfy the containment criterion relative to one sibling transcript.

**Figure 2.**
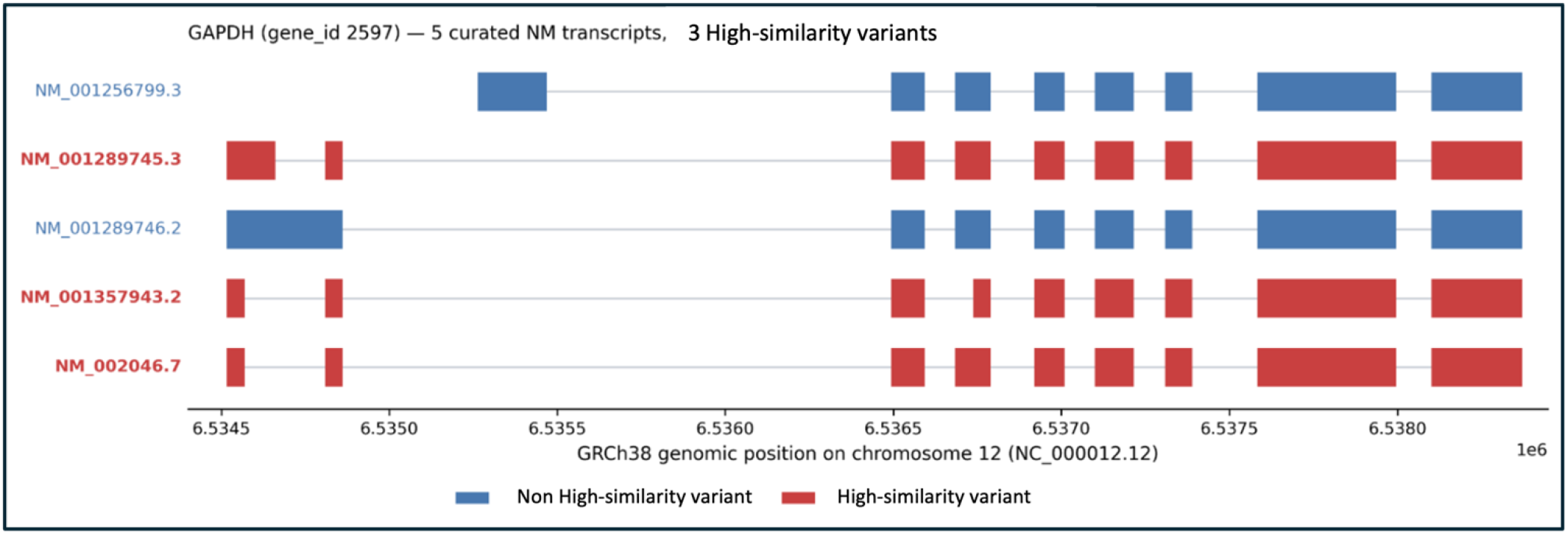
Exon-level structural comparison of curated GAPDH transcript variants illustrating high-similarity RNA variants. Exon structures of five curated *GAPDH* NM transcripts are shown according to GRCh38 genomic coordinates. Transcripts were classified as high-similarity RNA variants when every exon was fully contained within an exon of the same sibling NM transcript. Three transcripts - NM_001289745.3, NM_001357943.2, and NM_002046.7 (red) - met this criterion relative to NM_001289746.2, whereas NM_001256799.3 and NM_001289746.2 (blue) were not classified as high-similarity RNA variant.

Application of this criterion across the complete dataset is summarized in Figure 3. Of the 20,593 human protein-coding genes initially examined, 20,583 (99.95%) were successfully processed. Ten genes were excluded during transcript retrieval and filtering, including nine mitochondrial gene symbols for which corresponding records were not recovered and one ambiguous gene-symbol match. Among the processed genes, 962 (4.7%) lacked curated NM transcripts and were represented only by XM-predicted transcripts, and 7,805 (37.9%) had only a single curated NM transcript; these groups were therefore exempt from pairwise structural comparison. Within the mono-isoform set, 1,099 genes (5.3% of all processed genes) were single-exon genes, and 6,706 genes (32.6%) were multi-exon genes. The total multi-exon genes from mono-isoform and multi-isoform were 18,522 (90%). Notably, the proportion of single-exon mono-isoform genes is consistent with the ~6% of human genes reported to be single-exon in the alternative splicing analysis of Wang et al. (2008). By contrast, the much larger set of multi-exon genes represented here by only one curated NM transcript likely reflects the stringency of NCBI RefSeq curation and accession criteria, and therefore should not be interpreted as evidence that these genes lack alternative splicing, which has been estimated to affect more than 90% of human multi-exon genes. The remaining 11,816 genes with two or more curated NM transcripts were subjected to exon-wise containment analysis.

**Figure 3.**
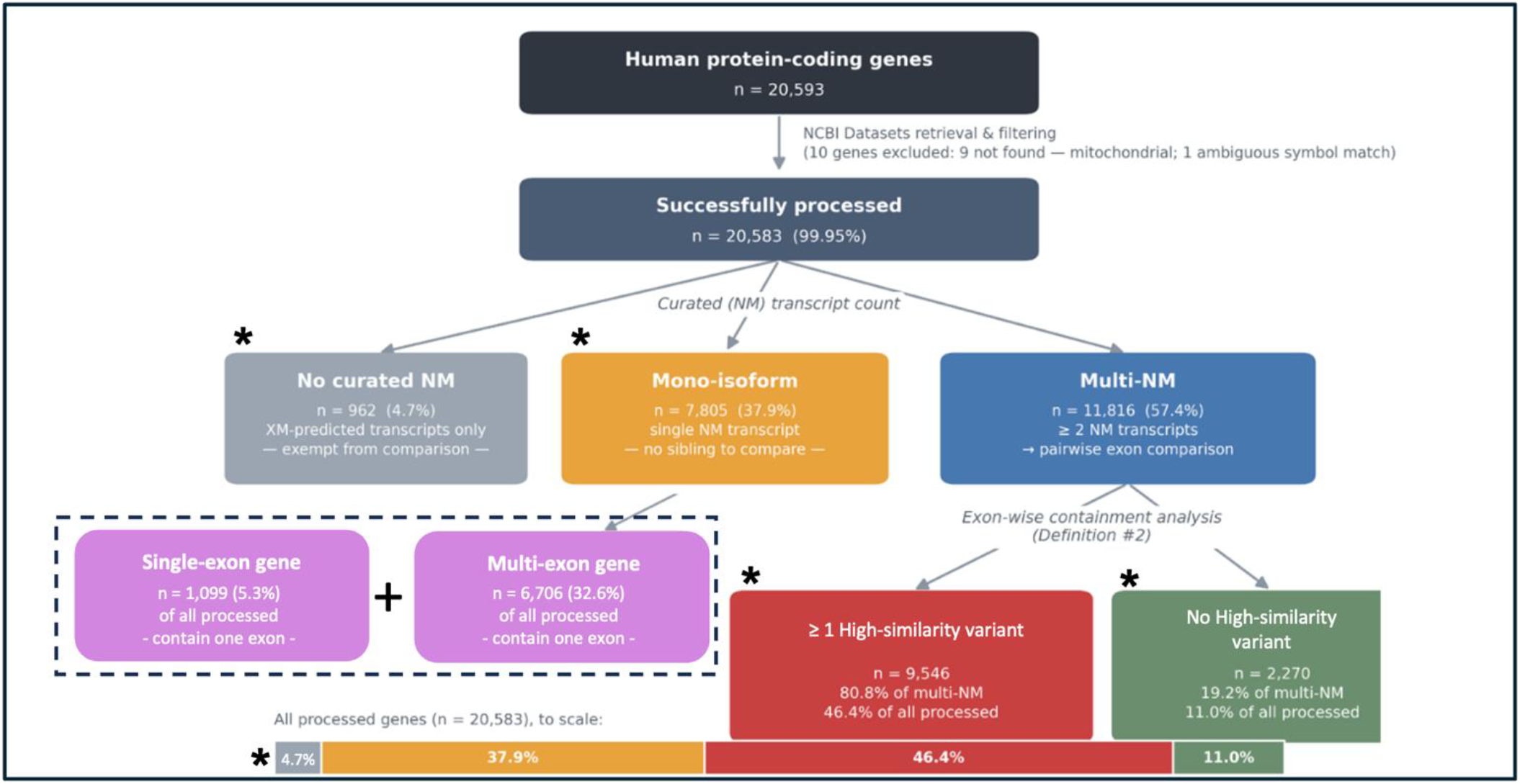
Distribution of curated RefSeq transcript isoforms and high-similarity variants across human protein-coding genes. A total of 20,593 human protein-coding genes were retrieved and filtered using NCBI Datasets, of which 20,583 (99.95%) were successfully processed. Based on the number of curated RefSeq NM transcripts, genes were classified as having no curated NM transcript (n = 962, 4.7%), a single NM isoform (n = 7,805, 37.9%), or multiple NM isoforms (n = 11,816, 57.4%). Among mono-isoform genes, 1,099 (5.3% of all processed genes) were single-exon genes and 6,706 (32.6%) were multi-exon genes. Multi-NM genes were further evaluated by pairwise exon-wise containment analysis to identify high-similarity transcript variants. Of these, 9,546 genes (80.8% of multi-NM genes; 46.4% of all processed genes) contained at least one high-similarity variant, whereas 2,270 (19.2% of multi-NM genes; 11.0% of all processed genes) did not. The proportional bar at the bottom summarizes the distribution of all 20,583 successfully processed genes.

Because structural containment alone does not determine whether a transcript can be selectively amplified, sequence-level discrimination must also be considered. A structurally overlapping transcript can still contain an informative nucleotide segment or exon-exon junction, whereas other variants may lack sufficiently distinctive exonic sequence and therefore require junction-level discrimination. We next evaluated this problem experimentally using representative high-similarity transcript variants.

### Exon–exon junction primer design enables discrimination of high-similarity HTRA1-AS1 variants

HTRA1-AS1 is a recently characterized long non-coding RNA (lncRNA) genetically associated with age-related macular degeneration (AMD) and has been implicated in AMD pathogenesis [22–24]. The HTRA1-AS1 locus contains multiple high-similarity RNA variants with extensively overlapping exon structures, providing a useful model for evaluating transcript variant–specific detection. To distinguish variants lacking sufficiently large transcript-specific exonic regions, we implemented an exon–exon junction (EEJ) primer-design strategy based on dual-sequence recognition across transcript-specific exon–exon junctions (Figure 4).

**Figure 4.**
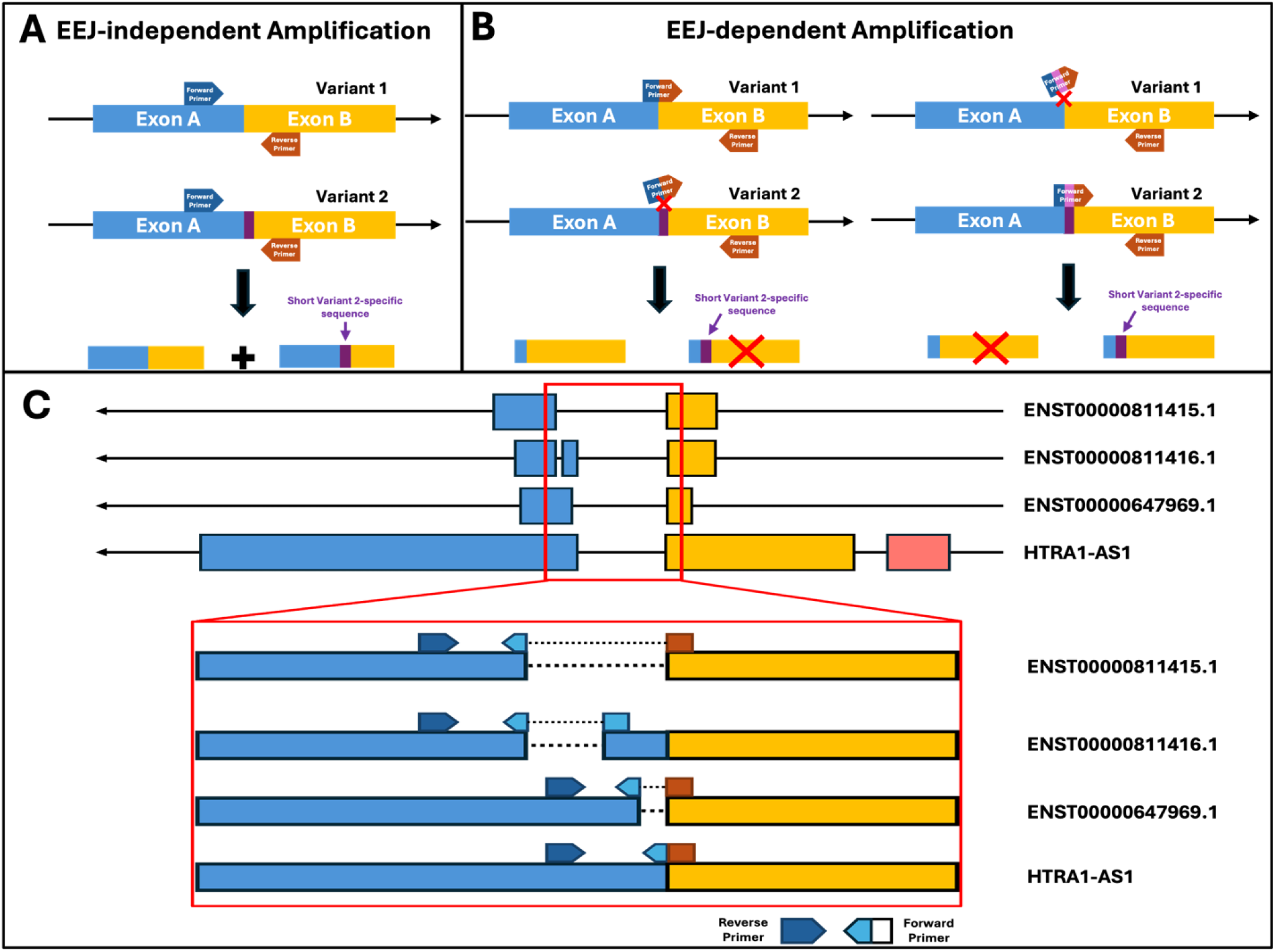
Exon–exon junction primers enable transcript variant–specific detection. **(A)** EEJ-independent amplification method: Primers designed within shared exons (Exon A and Exon B) amplify multiple transcript variants due to identical exon sequences, resulting in non-specific detection. **(B)** EEJ-dependent amplification method: A junction-spanning primer is designed across the exon–exon boundary. Amplification occurs only when the correct exon combination is present, enabling selective detection of the target variant, while mismatched variants are not amplified. **(C)** Application to *HTRA1-AS1* four transcript variants: Schematic of multiple variants lacking sufficiently distinguishable exon-internal regions. Junction-specific primer design targets distinct exon connectivity, allowing discrimination of closely related transcript variants. The inset illustrates primer positioning across exon–exon junctions for variant-specific amplification. Annotated accession identifiers for the HTRA1-AS1 transcripts (ENST0000811415.1, ENST0000811416.1, and ENST0000647969.1) were obtained from the Ensembl database.

In this strategy, one primer spans an exon–exon junction, with its 5′ portion complementary to the upstream exon and its 3′ portion complementary to the downstream exon. Efficient primer annealing therefore requires recognition of both exon sequences in the correct adjacency, providing junction-level specificity. A key design feature is the thermodynamic distinction between partial and complete primer binding. The Tm of each individual exon-recognizing arm was deliberately designed to be lower than the Tm of the full-length primer at least by >7 °C. Under these conditions, partial annealing to a single exon is thermodynamically unstable and was not detectably productive under the tested conditions. In contrast, when the primer encounters the correct exon–exon junction, both arms anneal cooperatively to form a stable duplex, enabling efficient amplification. This design suppresses amplification from transcripts sharing only a single exon as well as from genomic DNA containing intronic sequences.

The principle of this approach is illustrated in Figure 4A–B. Primers located entirely within shared exonic sequences cannot readily distinguish transcript variants containing identical target regions and may therefore amplify multiple variants (Figure 4A). In contrast, an EEJ-spanning primer recognizes the specific adjacency of two exons and preferentially amplifies transcripts containing the targeted exon–exon junction (Figure 4B). Transcript variants lacking the specified exon connectivity cannot support complete primer annealing, thereby providing a basis for discrimination among transcripts with otherwise highly similar exon structures.

We applied this strategy to the HTRA1-AS1 locus, in which multiple transcript variants exhibit extensive structural overlap but differ in their exon connectivity (Figure 4C). Because these variants provide limited variant-specific sequence within individual exons, EEJ-independent primers targeting shared exonic regions are insufficient for selective detection. We therefore designed EEJ primers targeting junctions characteristic of individual splice configurations. The inset in Figure 4C illustrates the positioning of these primers across selected exon–exon boundaries and demonstrates how differences in exon connectivity can be exploited for transcript variant–specific detection.

Using these design principles, we generated a panel of EEJ primers targeting distinct HTRA1-AS1 splice junctions. Four junction-specific forward primers were designed to recognize different exon–exon boundaries and were paired with a reverse primer. Amplification is therefore dependent on the presence of the targeted exon adjacency, providing a practical strategy for selectively detecting high-similarity RNA variants that cannot be readily distinguished using EEJ-independent primer design.

The calculated melting temperatures for each primer half and the full-length primers are summarized in Table 1, providing quantitative support for the thermodynamic basis of the design.

**Table 1.**
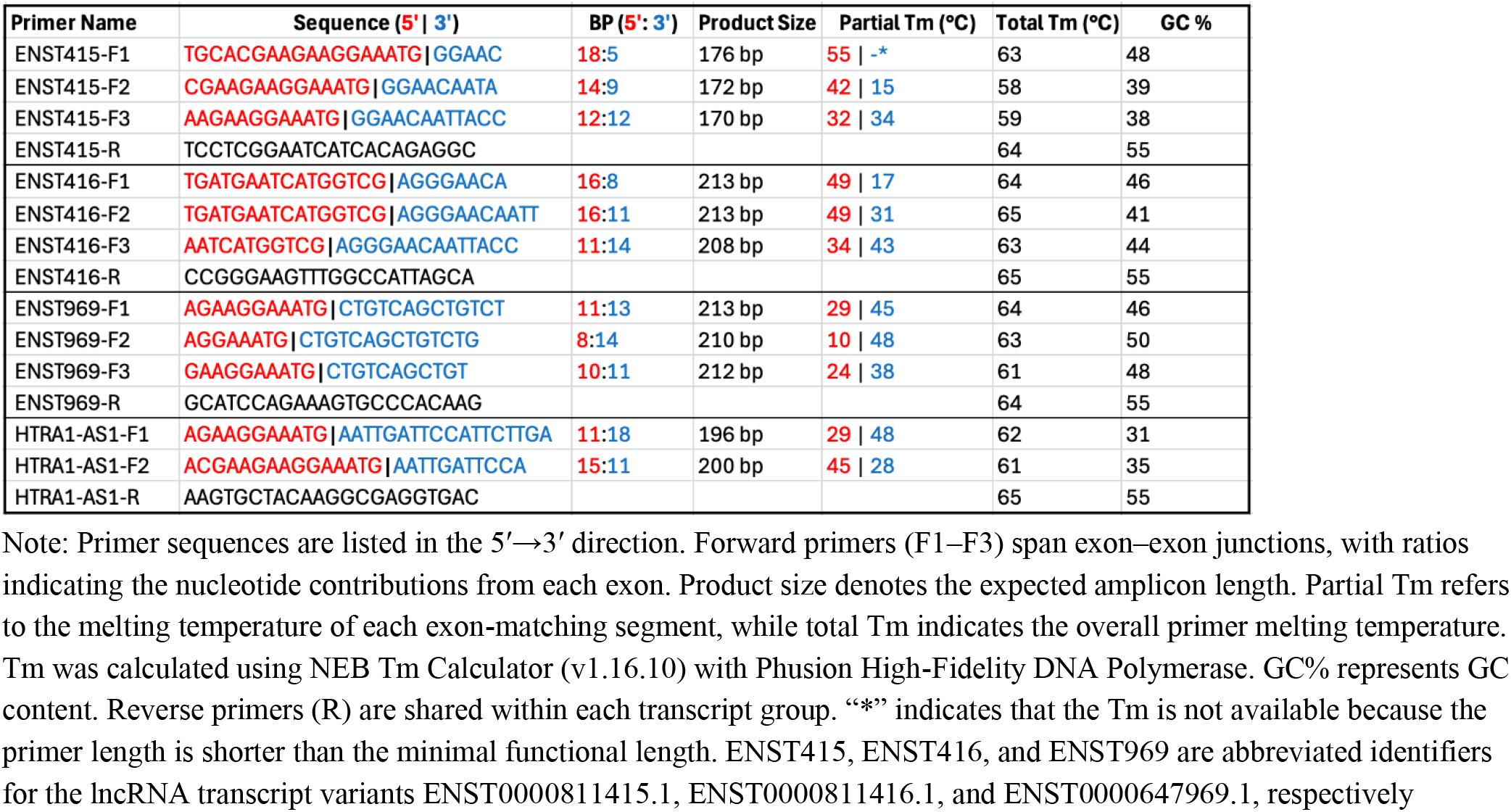
Exon–exon junction Primers and their Tm values.

### PCR optimization and sequence validation for HTRA1-AS1

PCR polymerase selection was critical for achieving robust and specific amplification using junction-spanning primers. We evaluated several commonly used PCR polymerases under identical cycling conditions and observed substantial differences in amplification efficiency and specificity. High-fidelity polymerases with optimized buffer systems consistently produced stronger and more reproducible products, whereas some enzymes yielded weak bands or exhibited nonspecific amplification. Notably, performance differences were evident even between similar Phusion High-Fidelity PCR Master Mix formulations obtained from different commercial suppliers. Based on these comparisons, Thermo Fisher Phusion Flash High-Fidelity PCR Master Mix and Qiagen 2× PyroMark PCR Master Mix (containing HotStarTaq DNA polymerase and 3 mM Mg^2+^) were selected for subsequent experiments. Both systems generated clear, specific bands with minimal background, demonstrating their suitability for exon–exon junction PCR (data for other polymerases not shown).

These findings underscore the importance of polymerase selection for the reliable detection of transcript variants lacking large distinguishable exonic regions. While predicted melting temperatures (Tm) provide a useful starting point for primer design and initial annealing temperature selection, the optimal annealing temperature must be determined empirically. Annealing conditions have a strong impact on assay performance and are also dependent on the PCR polymerase used. For example, the optimal annealing temperature for Phusion Flash High-Fidelity PCR Master Mix differs from that of PyroMark PCR Master Mix (Supplementary Data 1–3).

Using the optimized PCR conditions, each exon–exon junction primer set generated a single, discrete band at the expected size for its corresponding transcript variant (Figure 5A, 5C). No additional bands or smearing were observed, indicating high amplification specificity and minimal nonspecific products.

**Figure 5.**
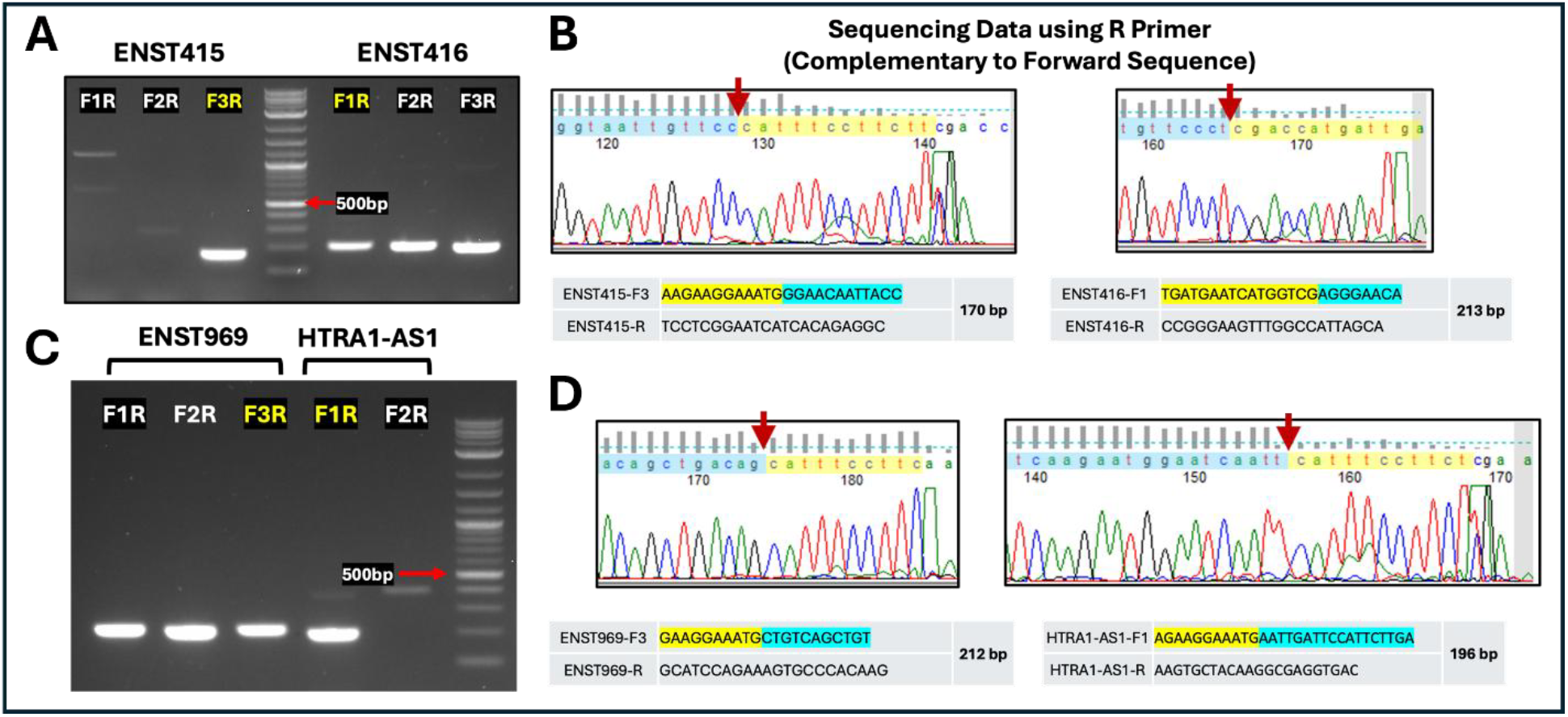
Variant-specific amplification of *HTRA1-AS1* transcripts using exon–exon junction primers. **(A, C)** Agarose gel electrophoresis showing PCR products generated with different primer combinations (F1R, F2R, F3R). Clear bands at the expected sizes (~170–213 bp) are observed for the correct variant, with minimal non-specific amplification. DNA ladders are included for size reference, with the ~500 bp band indicated. **(B, D)** Sanger sequencing confirms that the PCR products correspond to the expected exon–exon junction sequences. Red arrows mark the precise junction sites. Yellow and blue highlighted regions indicate the primer-binding sequences (including reverse-complement alignment), demonstrating accurate primer–template pairing and variant-specific amplification.

Importantly, primer sets targeting different splice junctions produced mutually exclusive amplification patterns across variants, consistent with junction-dependent recognition. The observed amplicon sizes (~170–213 bp) matched the predicted lengths based on primer design and exon connectivity, further supporting accurate target amplification.

To validate specificity at the sequence level, PCR products were directly subjected to Sanger sequencing without gel purification, providing a stringent test of amplification fidelity. Sequencing chromatograms obtained using reverse primers (complementary to the forward strand) exhibited clean, high-quality peaks with minimal background noise (Figure 5B, 5D). Critically, the sequences spanning the amplified regions precisely matched the designed exon–exon junctions, with the junction boundaries clearly identifiable at the expected positions (indicated by arrows). No evidence of mixed sequencing signals or off-target amplification was detected, indicating that a single dominant product was generated in each reaction.

For all tested transcript variants (ENST415, ENST416, ENST969, and *HTRA1-AS1*), the sequencing results confirmed that amplification occurred only when the correct exon adjacency was present, consistent with the double-recognition mechanism of the junction primers. Variants lacking the targeted exon–exon junction did not yield detectable products, demonstrating effective discrimination among closely related variants that share common exon sequences. Only the ENST415-specific forward primer F3, when paired with the common reverse primer (F3R), produced a specific PCR band, similar to the *HTRA1-AS1* forward primer F1 combined with the common reverse primer (F1R). In contrast, all three primer pairs performed efficiently for ENST416 and ENST969 under tested conditions.

### Empirical establishment of an arm-specific Tm constraint for EEJ primer design

To establish a thermodynamic criterion for minimizing productive amplification from partial EEJ-primer binding, we examined the relationship between primer melting temperature (Tm) and PCR amplification at a fixed annealing temperature. Eleven primers (A–K) with progressively shortened HTRA1-AS1-complementary sequences were paired with a common reverse primer and used to amplify HTRA1-AS1 from human retinal cDNA (Figure 6). The primers were designed to produce progressively lower Tm values, resulting in differences between the fixed annealing temperature (Ta = 62 °C) and primer Tm (Ta − Tm) ranging from 0 to 14°C.

**Figure 6.**
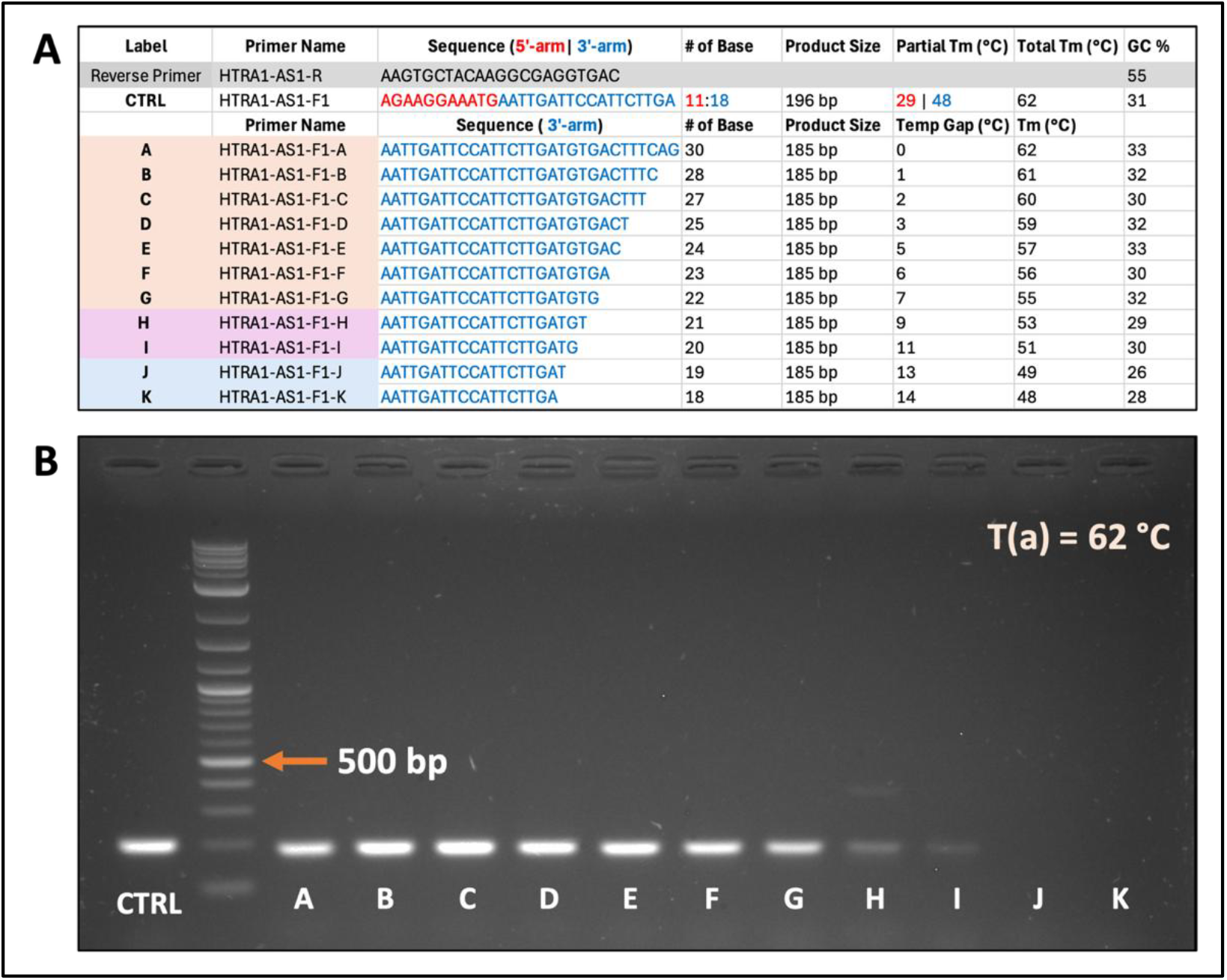
Empirical evaluation of the relationship between primer Tm and amplification efficiency. **(A)** Primer sequences and calculated Tm values for a series of HTRA1-AS1 primers containing progressively shortened shared sequences. The difference between the annealing temperature (Ta = 62°C) and primer Tm ranged from 0 to 14°C. **(B)** Gel electrophoresis of PCR products generated using the corresponding primer series and a common partner primer. Primers A–G, with Tm differences of 0–7°C, produced amplification comparable to the EEJ control primer (CTRL). Primers H and I, with differences of 9 and 11°C, respectively, showed reduced but detectable amplification, whereas primers J and K, with differences of 13 and 14°C, produced no detectable PCR products under the tested conditions.

Primers A–G, with Ta − Tm differences of 0–7°C, generated PCR products comparable in intensity to the control EEJ primer (CTRL). Amplification decreased but remained detectable with primers H and I, corresponding to differences of 9 and 11°C, respectively. In contrast, primers J and K, with differences of 13 and 14°C, produced no detectable amplification under the same conditions. These results demonstrate a progressive loss of amplification efficiency as primer Tm decreases relative to Ta, with amplification becoming undetectable at a Ta − Tm difference of approximately 13–14°C under the tested conditions.

We next examined whether a noncomplementary 5′ sequence could permit amplification when only the 3′ arm of an EEJ primer was correctly matched to the template. Six primers were generated in which the same HTRA1-AS1-complementary 3′ arm, with a calculated Tm of 48°C, was joined to random 5′ sequences of identical length (Figure 7). The random sequences were generated using the Random Sequence Generator at Bioinformatics.org. The resulting full-length primers had calculated Tm values within approximately ±3 °C of the control EEJ primer, thereby maintaining similar overall primer thermodynamic properties while disrupting sequence-specific recognition by the 5′ arm. Each primer was tested across annealing temperatures from 58 to 64°C.

**Figure 7.**
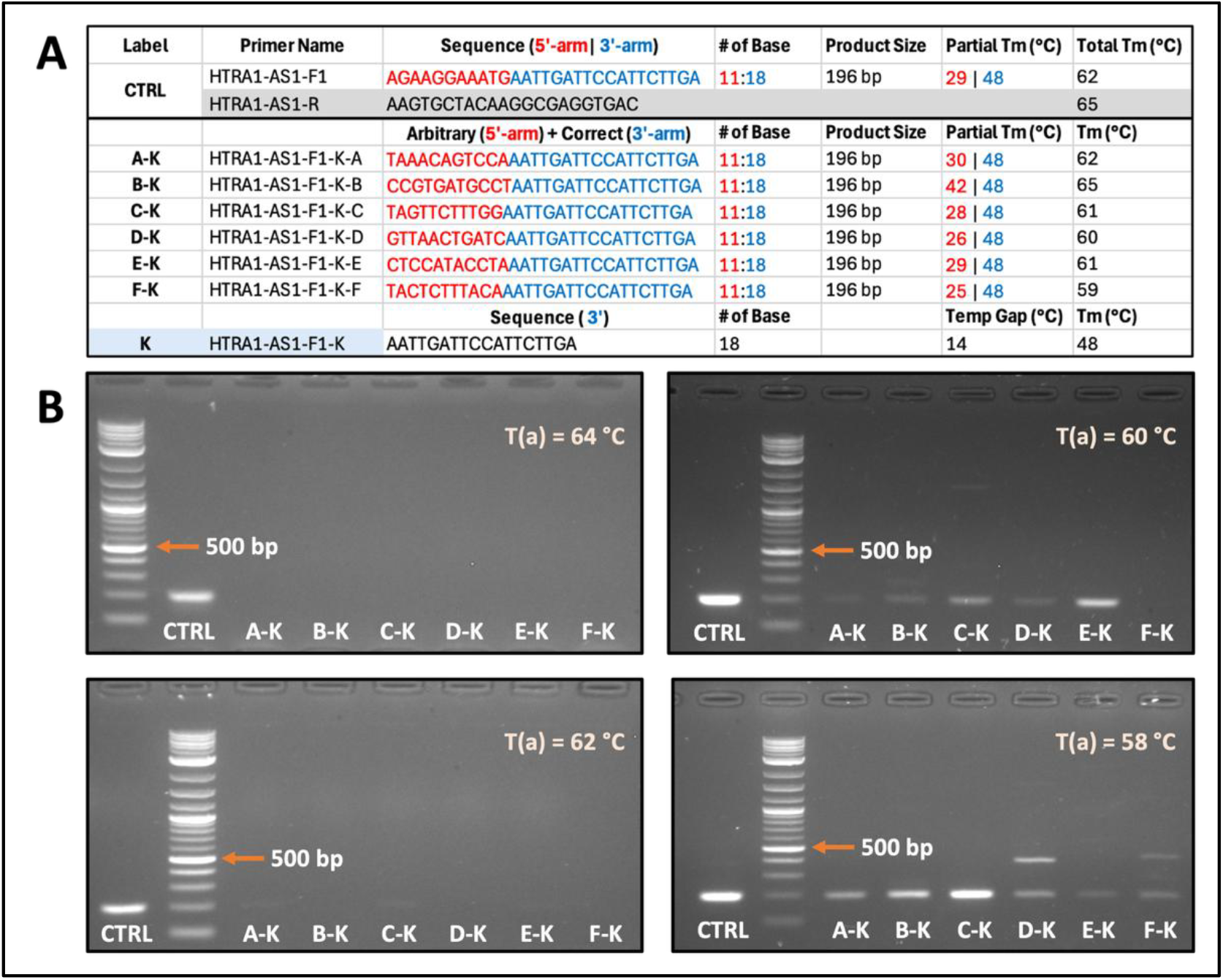
Effect of a noncomplementary 5′ arm on amplification mediated by a shared 3′ primer segment. **(A)** Sequences and calculated Tm values of six primers containing an identical HTRA1-AS1-complementary 3′ arm (Tm = 48°C) joined to random, noncomplementary 5′ sequences of equal length. Full-length primer Tm values were maintained within approximately ±3 °C of the control EEJ primer. **(B)** PCR amplification performed at annealing temperatures of 58, 60, 62, and 64°C. Amplification from primers containing random 5′ arms decreased as the annealing temperature increased and was undetectable at 64°C, whereas the correctly matched EEJ control primer remained functional. These results indicate that a correctly matched 3′ arm alone becomes insufficient to support detectable amplification when its Tm is approximately 15°C or more below the annealing temperature.

At lower annealing temperatures, amplification was observed with several primers containing random 5′ arms, indicating that the correctly matched 3′ segment alone could support extension when the random 5′ sequence provided sufficient incidental stabilization. Amplification progressively decreased as the annealing temperature increased. At Ta = 64°C, none of the primers containing a random 5′ arm produced a detectable PCR product by agarose gel electrophoresis, whereas the correctly matched EEJ control primer remained efficiently amplified. Because the shared 3′ arm had a Tm of 48°C, this condition corresponded to a separation of approximately 16°C between the arm Tm and the annealing temperature. These findings indicate that when the Tm of the correctly matched 3′ segment is sufficiently below the annealing temperature, binding of that segment alone is thermodynamically insufficient to support detectable amplification, even when incorporated into a full-length oligonucleotide with an overall Tm comparable to that of a functional EEJ primer.

Taken together, the sequential-Tm experiment demonstrated a marked loss of amplification when primer Tm was approximately 13–14°C below the annealing temperature, while the random-arm experiment showed complete suppression of detectable amplification when the correctly matched 3′ segment was approximately 16°C below the annealing temperature. Based on these complementary observations, we adopted a 15°C Tm-gap as a conservative design constraint for EEJ primers, such that the Tm of each individual exon-matching arm is designed to be at least 15°C below the PCR annealing temperature.

This criterion is intended to minimize productive extension from either arm independently while retaining stable annealing of the complete primer across the intended exon–exon junction. Primers designed according to this constraint were subsequently generated and experimentally evaluated to further validate the EEJ-primer design strategy.

### Tm-guided EEJ primer design supports transcript-specific quantitative PCR

To determine whether the experimentally derived Tm-guided design rule could be implemented automatically, transcript-specific primer pairs for APEX1 and GAPDH were generated using TransPrimer and experimentally evaluated alongside the HTRA1-AS1 primer sets. Thus, the APEX1 and GAPDH assays served as experimental validation of the automated TransPrimer workflow, rather than of manually optimized junction-primer designs.

To evaluate the amplification specificity and quantitative performance of Tm-guided exon–exon junction (EEJ) primers, real-time quantitative PCR (qPCR) assays were performed for 12 transcript variants from three human genes (APEX1, HTRA1-AS1, and GAPDH). High-resolution melting-curve analysis using HEK293T cell cDNA revealed single, sharp melting peaks for all evaluated transcripts, with no evidence of nonspecific amplification or primer-dimer formation (Figure 8). Single-peak profiles were observed for APEX1 variants NM_001641.4 (NM641), NM_001244249.2 (NM249), NM_080648.3 (NM648), and NM_080649.3 (NM649); HTRA1-AS1 isoforms ENST415, ENST416, ENST969, and HTRA1-AS1; and GAPDH variants NM_002046.7 (NM046), NM_001289745.3 (NM745), NM_001256799.3 (NM799), and NM_001357943.2 (NM943). For amplification of GAPDH NM_002046.7, the non-coding RNA variant NR_152150.2 is co-amplified, yielding an amplicon of identical size and sequence because NM_002046.7 cannot be selectively amplified independently of NR_152150.2. Alternative primer designs that exclude NR_152150.2 would require targeting regions shared with additional GAPDH transcripts, resulting in broader co-amplification. Both single-EEJ (NM046-A) and dual-EEJ (NM046-B) primer-pairing strategies amplified the intended transcripts with comparable specificity. Single size products were further confirmed by agarose gel electrophoresis (Supplementary Data 4). These results support the ability of the Tm-guided strategy to amplify closely related transcript variants while minimizing detectable nonspecific products and primer-dimer formation.

**Figure 8.**
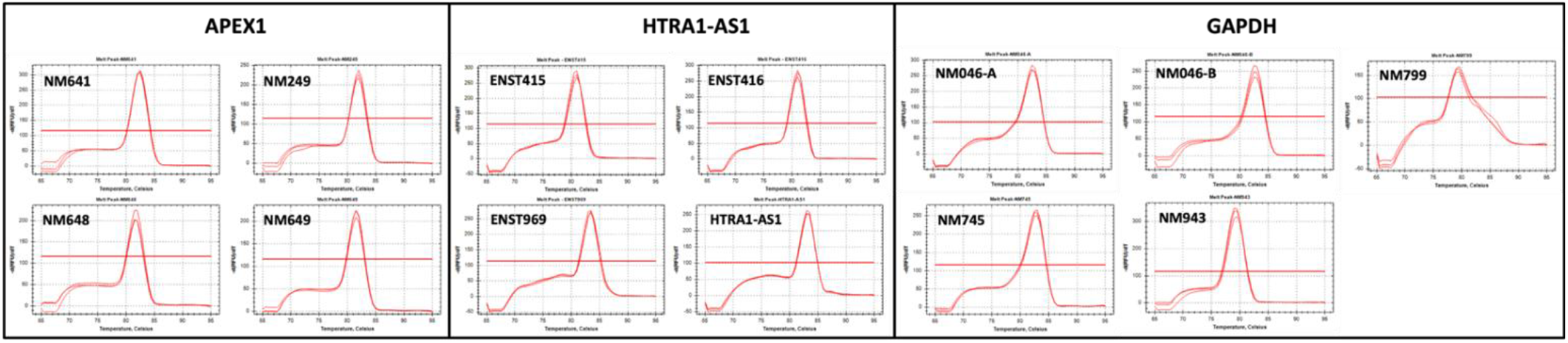
Melting curve analysis of Tm-guided exon–exon junction (EEJ) primers across target transcript variants. Real-time qPCR dissociation curves for 12 transcript variants across three human genes (APEX1, HTRA1-AS1, and GAPDH) amplified from HEK293T cDNA (8.3 ng input total RNA equivalent). Single, sharp melting peaks confirm target-specific amplification without off-target products or primer-dimers. NM046-A and NM046-B denote single- and dual-EEJ primer pairs targeting GAPDH NM_002046.7 variant along with NR_152150.2. Reactions were performed in triplicate using iTaq Universal SYBR Green Supermix on a Bio-Rad CFX384 system.

The quantitative performance and amplification linearity of the designed EEJ primers were further characterized using a five-point, 10-fold serial dilution series ranging from 16.6 ng to 1.66 pg of input-RNA equivalent per reaction (Figure 9). Most primer pairs exhibited amplification efficiencies within the commonly accepted 90–110% range with R^2^ > 0.98 across a four-log dynamic range. For example, APEX1 NM641 yielded an efficiency of 95.1% (R^2^ = 0.997), whereas GAPDH NM046-B yielded 97.6% (R^2^ = 0.999). At the 10,000-fold dilution point (approximately 2 pg input-RNA equivalent), increased Cq variance and curve dispersion were observed for lower-abundance isoforms, consistent with stochastic sampling at low template abundance. A complete standard curve could not be generated at the highest dilutions for the low-abundance GAPDH variant NM746, although single-peak amplification was retained at standard cDNA inputs. Together, these results indicate that the Tm-guided EEJ design is compatible with robust qPCR kinetics across a broad range of template concentrations.

**Figure 9.**
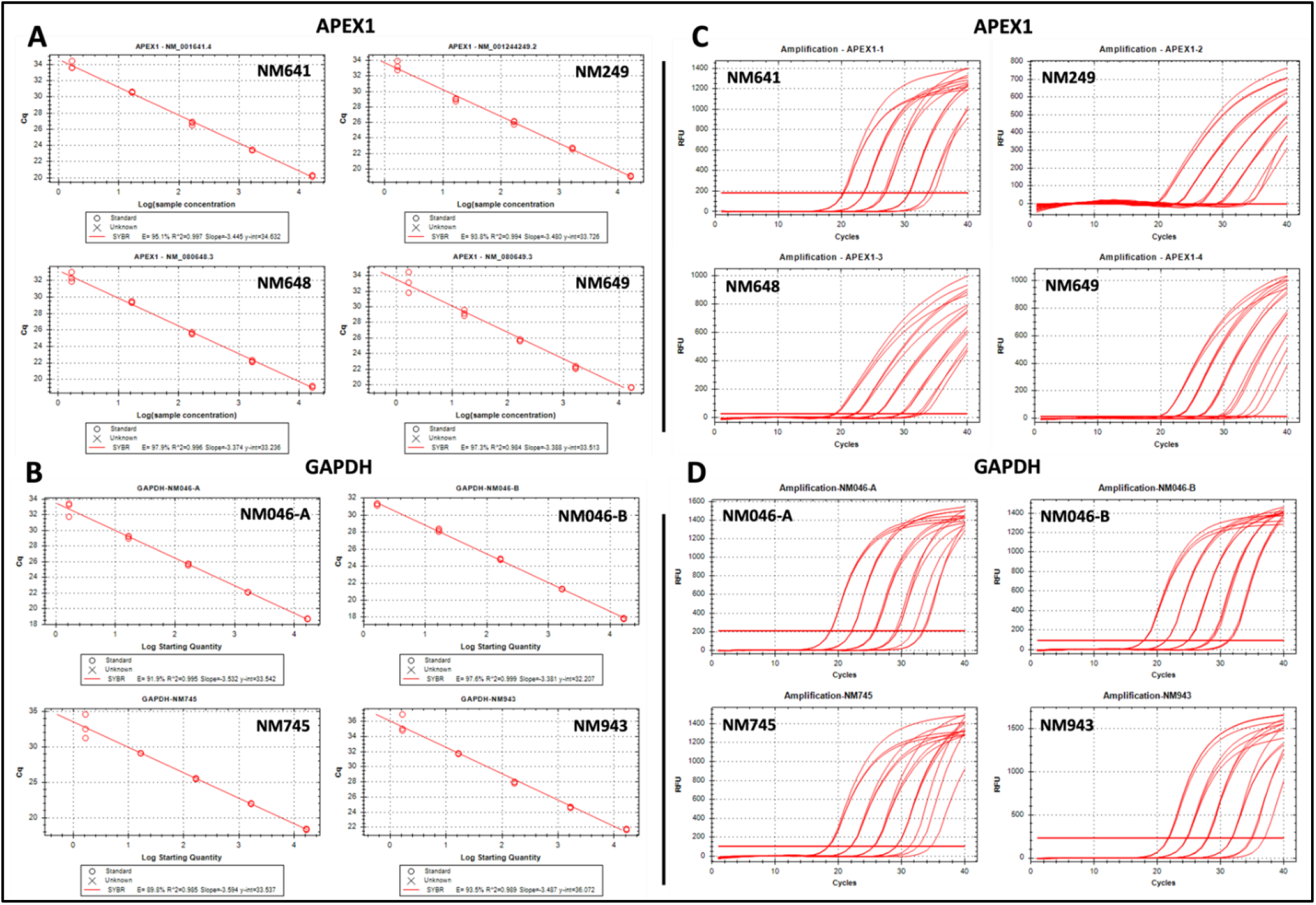
Quantitative efficiency and amplification kinetics of Tm-guided EEJ primers across serial sample dilutions. Standard curves (A, B) and representative real-time amplification plots (C, D) derived from five-point, 10-fold serial dilutions of HEK293T cDNA (ranging from 16.6 ng down to 1.66 pg input RNA equivalent per reaction). Standard curve linear regression demonstrates high amplification efficiency (E = 89.8% - 97.9%) and high coefficient of determination (R^2^ > 0.98) across tested APEX1 and GAPDH transcript variants over a 4-log dynamic range.

### Exon–exon junction RT-PCR reduces heteroduplex formation during simultaneous detection of HTRA1-AS1 transcript variants

Four HTRA1-AS1 variants (ENST415, ENST416, ENST969, and HTRA1-AS1) were initially coamplified using a common primer set to assess amplification specificity. Agarose gel electrophoresis revealed three bands corresponding to ENST416 (305bp), HTRA1-AS1 (481bp), and an unexpected intermediate band (about 400bp) between them (Figure 10A). Sanger sequencing of the gel-extracted intermediate band confirmed that it represented a heteroduplex composed of hybrid strands derived from ENST416 and HTRA1-AS1 (Figure 10C).

**Figure 10.**
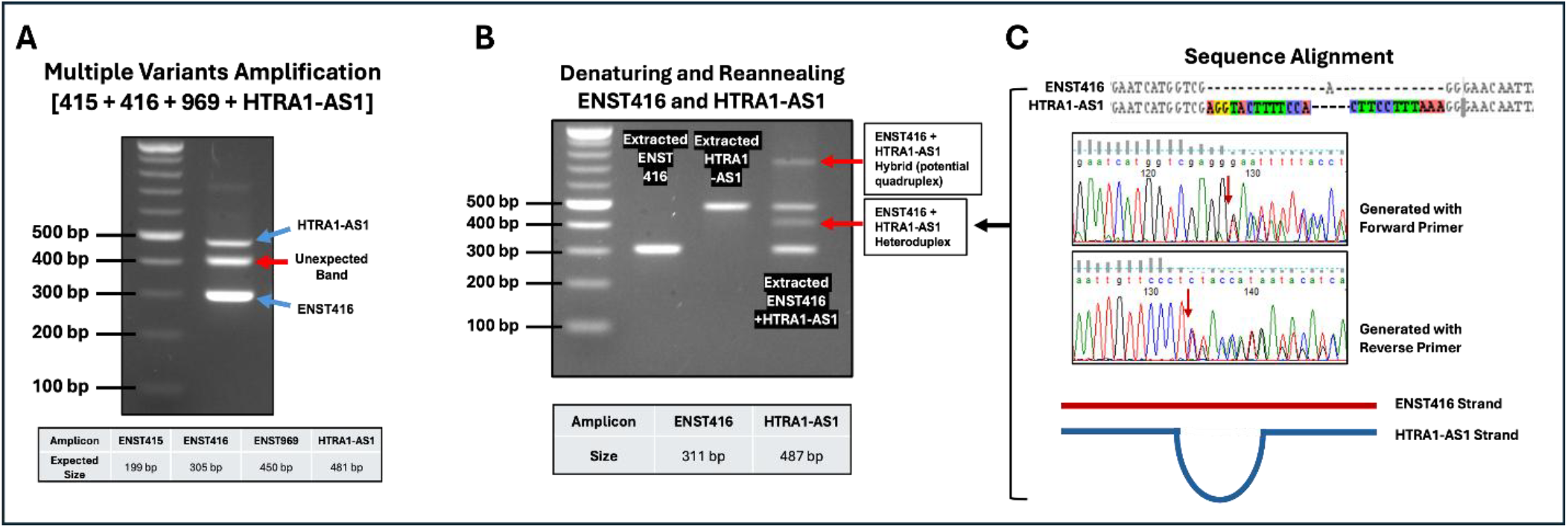
Heteroduplex and quadruplex formation during detection of RNA variants using EEJ-independent RT-PCR. (**A**) Co-amplification of four HTRA1-AS1 variants (ENST415, ENST416, ENST969, and HTRA1-AS1) using a common primer set produces multiple bands on agarose gel electrophoresis, including expected products and an unexpected intermediate band (red arrow), suggesting nonspecific hybridization. (**B**) Validation of heteroduplex formation by denaturation and reannealing. After denaturation and reannealing single bands together, which individually ENST416 and HTRA1-AS1 products, additional higher-molecular-weight bands (red arrows) were generated, consistent with heteroduplex formation between the two variants. Amplicon sizes are indicated below. (**C**) Sequence alignment and Sanger sequencing traces of the heteroduplex product generated using forward and reverse primers. Overlapping chromatogram peaks and misaligned base calls confirm the presence of hybrid DNA strands derived from ENST416 and HTRA1-AS1. A schematic model illustrates heteroduplex formation through partial base pairing between homologous regions of the two transcripts.

To exclude the possibility that this band resulted from co-migrating independent amplicons, gel purified ENST416 and HTRA1-AS1 PCR products were subjected to denaturation (98 °C) followed by reannealing (63 °C). Individual samples produced a single band, whereas the mixed sample generated an additional band (~400 bp), consistent with heteroduplex formation between the two variants and even a higher molecular weight higher-order heteromeric band (Figure 10B).

As illustrated in Figure 11, partial sequence homology between amplicons of different lengths can contribute to the formation of proposed heteroduplex and higher-order structures (quadruplex) during reannealing. PCR products or these intermediates can further assemble into higher-order structures, including loop-containing quadruplexes and multi-stranded quadruplexes, which contribute to aberrant bands observed on agarose gels. The presence of unpaired regions and overlapping homologous segments facilitates these interactions, highlighting how shared sequence regions between PCR products drive heteroduplex and higher-order complex formation.

**Figure 11.**
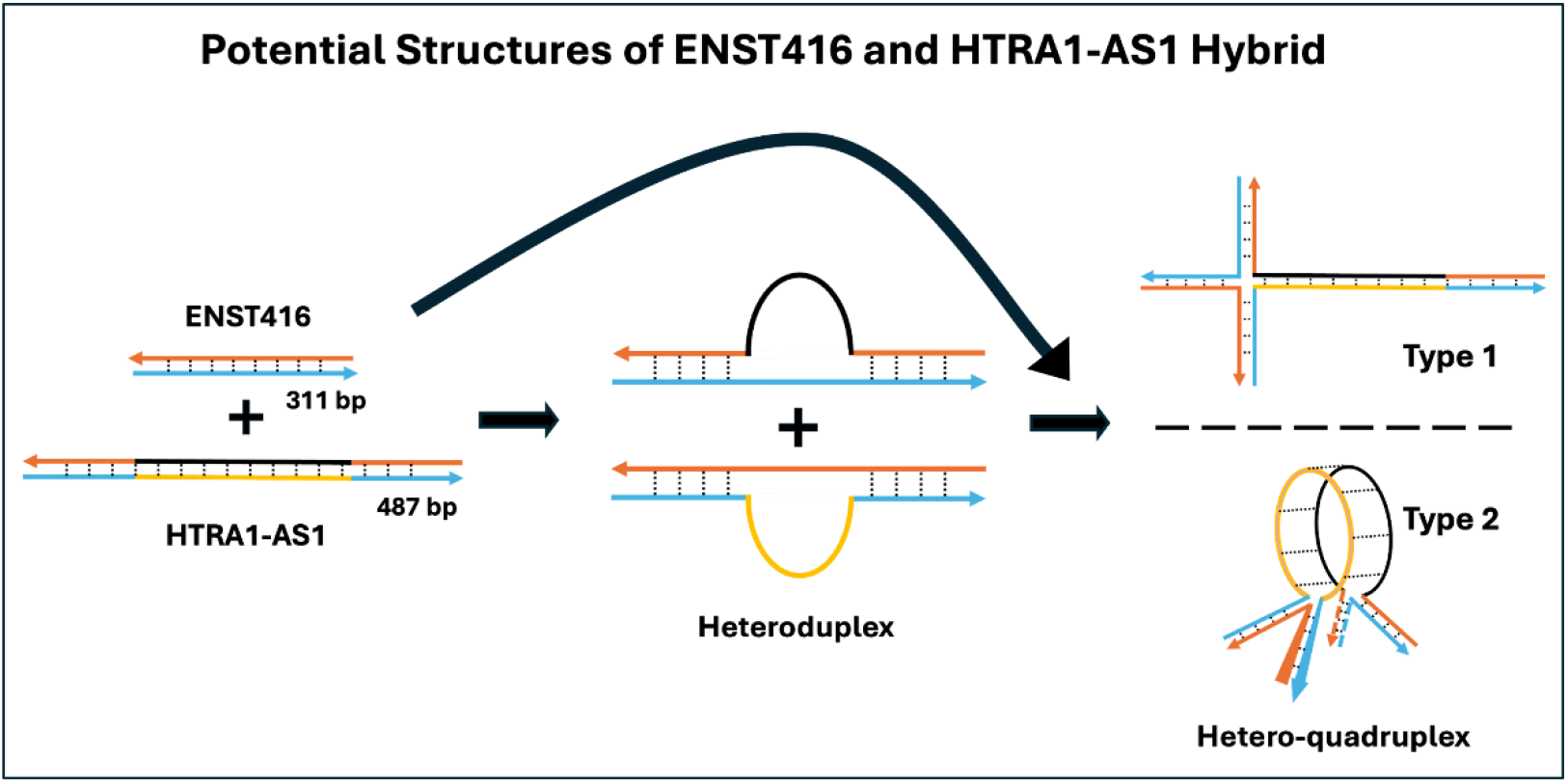
Potential hybrid structures formed between ENST416 and HTRA1-AS1 PCR products. Schematic illustrating the formation of hybrid DNA structures between ENST416 (311 bp) and HTRA1-AS1 (487 bp) amplicons containing partially homologous regions. Upon denaturation and reannealing, the shared sequences enable intermolecular base pairing, leading to heteroduplex formation with single-stranded loop regions corresponding to non-homologous segments. Two possible higher-order structures are proposed: (Type 1) a cruciform-like heteroduplex formed through crosswise pairing of homologous regions, and (Type 2) a higher-order heteromeric-like assembly involving multistrand interactions and circularized pairing of complementary segments. Arrows indicate strand orientation, and dashed lines represent base-paired regions. These structures may contribute to heteroduplex DNA artifacts observed during PCR amplification.

We next evaluated strategies to reduce heteroduplex or higher-order complex formation. An additional primer set (P1) was used to selectively co-amplify ENST416 and HTRA1-AS1 variants, resulting in the formation of heteroduplex products alongside the two variant amplicons. To minimize extensive homologous regions, a second primer configuration (P2) was designed using a common forward primer and two distinct reverse primers—including an exon–exon junction primer—that uniquely anneal to each variant (Supplementary Data 2). This design excludes the potential for loop formation between variants. Heteroduplex band was still observed with the P2 primer combination; however, its intensity was reduced (Figure 12).

**Figure 12.**
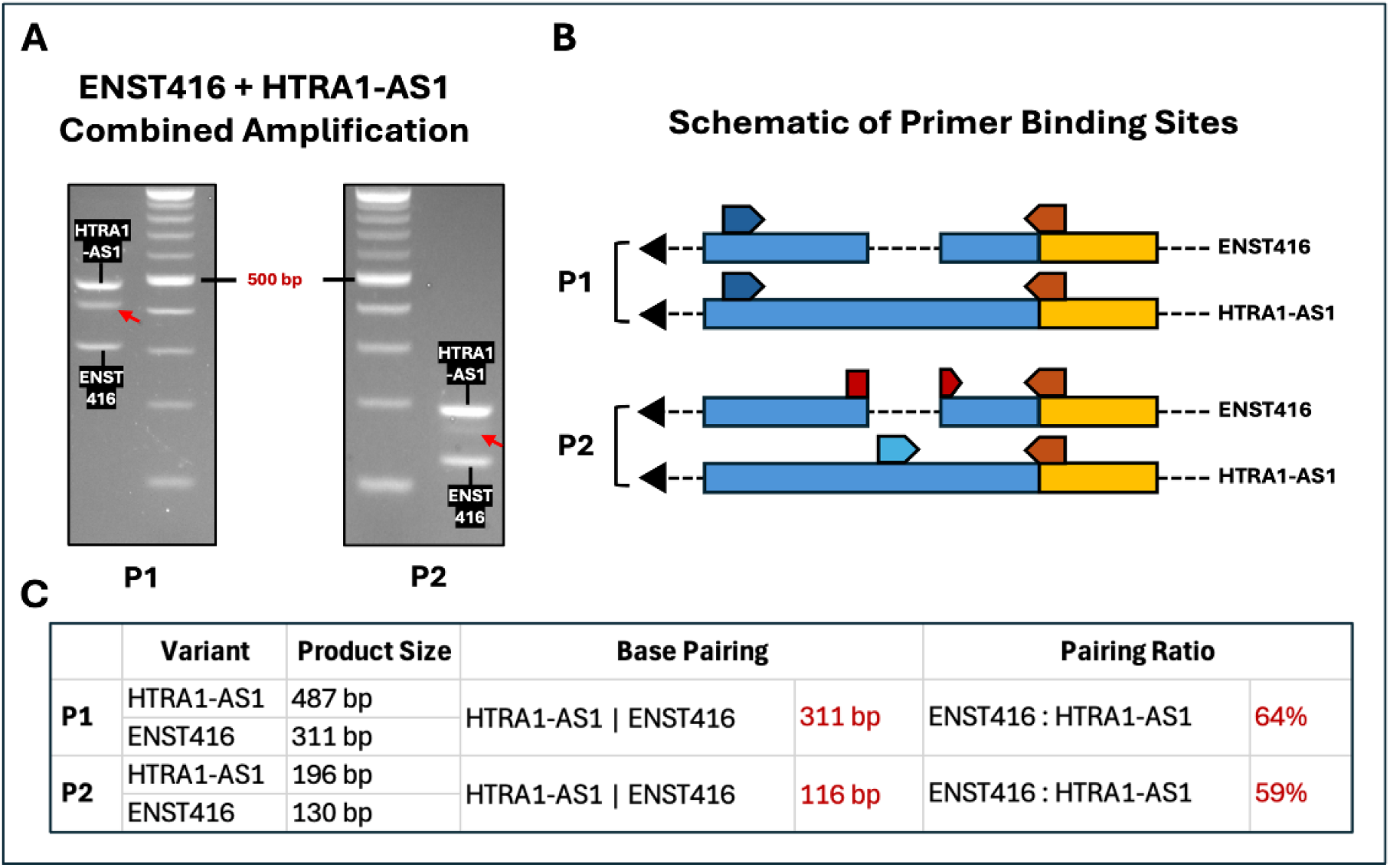
Exon–exon junction RT-PCR reduces heteroduplex formation. (**A**) Agarose gel electrophoresis showing co-amplification of ENST416 and HTRA1-AS1 using different primer designs. Exon-spanning primer set (P1) generates strong heteroduplex formation, evident as an additional band between the expected amplicons. Use of variant-specific reverse primers (P2) reduces heteroduplex formation, although intermediate bands remain detectable. (**B**) Schematic representation of primer design strategies and their effects on heteroduplex formation. P1 employs shared primer binding regions, promoting hybridization between partially homologous amplicons. P2 introduces variant-specific primers to reduce overlap. The reduced sequence complementarity and junction-specific targeting together minimize heteroduplex formation.

Thus, exon–exon junction RT-PCR provides greater flexibility in primer design, enabling reduction of heteroduplex formation by limiting amplification of homologous regions between variants (Figure 10A–C).

Collectively, these results demonstrate that heteroduplex formation in EEJ-independent RT-PCR can arise from partial sequence complementarity between coamplified variants and can be substantially minimized by exon–exon junction–targeted primer design. The EEJ-dependent strategy enables highly specific, sequence-validated detection of transcript variants, even when the exon-spanning primer approaches fail to resolve closely related splice isoforms.

## Discussion

Alternative transcript usage is a pervasive feature of the human transcriptome, and previous genome-wide studies have established extensive isoform diversity at individual gene loci. EST-based analyses demonstrated widespread alternative isoform production, while ENCODE transcriptome profiling showed that genes frequently express multiple isoforms and that expressed isoform diversity increases with the number of annotated transcripts [25, 26]. More recent long-read transcriptome analyses and continued refinement of GENCODE annotations have further revealed the complexity of human transcript structures [27, 28]. Our analysis extends these observations by addressing a distinct and experimentally important dimension of transcript diversity: the structural similarity among transcript variants derived from the same gene. Among 20,583 human protein-coding genes analyzed, 7,805 (37.9%) contained a single curated RefSeq NM transcript, whereas 11,816 (57.4%) contained multiple NM transcripts (Figures 1 and 3). Strikingly, 9,546 of these 11,816 multi-transcript genes (80.8%), corresponding to 46.4% of all analyzed protein-coding genes, contained at least one high-similarity transcript variant (Figure 3). Moreover, the prevalence of high-similarity variants increased markedly with transcript multiplicity, from approximately 60% among genes with two NM transcripts to >80% among genes with three transcripts, >90% among genes with four or five transcripts, and nearly all genes with larger transcript repertoires (Figure 1). These findings demonstrate that transcript number alone does not fully capture the practical complexity of transcript-specific detection; rather, the structural relationships among sibling transcripts are critical determinants of whether individual variants can be selectively detected. Among the 7,805 genes represented by a single curated NM transcript, 1,099 were single-exon genes and 6,706 were multi-exon genes (Figure 3). The substantial representation of single-exon genes is consistent with previous genome-wide surveys documenting a distinct population of human single-exon or intronless protein-coding genes [25, 29–31], although reported numbers vary with genome annotation and gene-definition criteria. Importantly, the much larger group of 6,706 multi-exon genes represented by only a single curated NM transcript demonstrates that the structural potential for alternative splicing does not necessarily translate into multiple curated transcript isoforms. This finding also suggests that NCBI-curated NM transcripts represent a more selective transcript set than those available from other transcript annotation sources. Together, our genome-wide analysis identifies high transcript similarity as a widespread feature of the curated human transcriptome and a major challenge for transcript-specific RT-PCR/qPCR. Because high-similarity variants may share extensive exonic sequence and differ primarily in exon connectivity, these findings provide a quantitative rationale for exploiting transcript-specific exon–exon junctions as discriminatory sequence features for primer design.

Accurate detection of transcript variants is important for studies of alternative splicing, transcript-specific regulation, and isoform-dependent biological function. Although sequencing technologies can resolve transcript diversity at scale, targeted RT-PCR and qPCR remain widely used for validation and quantitative analysis. These assays become particularly difficult when related transcripts share extensive exonic sequence and differ primarily in the connectivity of otherwise common exons. Our genome-scale structural analysis illustrates the extent of this challenge within curated RefSeq annotations: 9,546 protein-coding genes contained at least one high-similarity NM transcript under the exon-containment criterion used here. This classification does not imply that every such transcript requires an EEJ primer; rather, it identifies a large set of genes in which EEJ-independent exon-specific design may be constrained and junction-level information may provide additional discriminatory power.

The central methodological contribution of this study is an experimentally defined thermodynamic rule for EEJ-primer design. Rather than specifying only a fixed number of nucleotides on each side of a splice junction, the method evaluates the 5′ and 3′ exon-matching arms independently relative to the PCR annealing temperature. In the sequential-Tm experiment, amplification declined as primer Tm fell below Ta and became undetectable at a Ta−Tm difference of approximately 13–14 °C. In the complementary random-arm experiment, a correctly matched 3′ arm with a Tm approximately 16 °C below Ta did not support detectable amplification when the 5′ arm was noncomplementary, whereas the fully matched junction primer remained functional. On the basis of these complementary observations, we adopted an approximately 15 °C arm-specific Ta−Tm gap as a conservative design criterion. This experimentally derived rule provides a quantitative basis for limiting productive extension from partial EEJ-primer binding while preserving stable full-length binding across the intended junction.

Experimental testing supported the specificity of this approach across several transcript contexts. Junction-directed primers distinguished closely related HTRA1-AS1 variants, and Sanger sequencing confirmed that the amplified products contained the intended exon–exon junctions. In qPCR experiments spanning 12 transcript variants from HTRA1-AS1, APEX1, and GAPDH, the assays generated single melting peaks without detectable secondary products under the conditions tested. Most primer pairs also showed amplification efficiencies within the commonly accepted 90–110% range and R^2^ values above 0.98 across a four-log dilution series. These results indicate that restricting primer placement to exon junctions while applying arm-specific Tm criteria can preserve quantitative performance as well as specificity.

The quantitative PCR experiments targeting APEX1 and GAPDH directly tested whether the experimentally derived Tm-guided design principle could be incorporated into an automated transcript-specific primer-design workflow. TransPrimer integrates the key steps required for this process, including transcript comparison, identification of discriminating EEJs, independent evaluation of full-length primer and 5′/3′ arm Tm values, and generation of compatible primer pairs. The single-peak amplification and robust quantitative performance of the TransPrimer-generated assays support the feasibility of translating the arm-specific Tm criterion into an automated design strategy. These results therefore extend the experimentally established Tm-guided principle beyond manually optimized junction primers and demonstrate its practical application to automated transcript-specific primer design.

The experiments with HTRA1-AS1 also highlight a practical problem that can arise when highly homologous transcript variants are coamplified. Products sharing extensive sequence can reanneal to form heteroduplex species that migrate anomalously on agarose gels and complicate band interpretation. Sanger sequencing and denaturation–reannealing experiments supported the formation of such heteroduplex products in our system. Junction-directed primer configurations reduced these aberrant products by limiting the extent of shared amplicon sequence and increasing variant selectivity. We therefore view reduced heteroduplex formation as an additional practical benefit of transcript-specific primer design rather than as the primary mechanistic basis of the method.

Several existing computational resources provide complementary capabilities for exon-exon junction or isoform-aware primer design [15–17]. Primer-BLAST can require a primer to span an exon junction and allows users to specify the minimum number of matching nucleotides on the 5’ and 3’ sides of the junction, while emphasizing broad sequence-specificity assessment [17]. Ex-Ex Primer was developed specifically for junction-spanning oligonucleotides and includes a threshold for the Tm difference between the complete junction primer and a partially matching sequence; the published threshold was modified on the basis of an observation made during initial experimental validation rather than derived from a systematic Tm-versus-amplification series [16]. ExonSurfer provides an integrated qPCR workflow with automatic selection of informative junctions, transcript/off-target analysis, SNP avoidance, and user-adjustable primer length, GC content, Tm, and junction-overlap parameters [15].

These developments underscore the need for transcript-aware PCR design, while the present study addresses a distinct experimental question: how far the Tm of each partially matched EEJ-primer arm must fall below the annealing temperature before productive amplification is suppressed.

Several limitations should be considered. The genome-wide structural analysis was restricted to curated RefSeq NM transcripts mapped to GRCh38 and therefore does not encompass all experimentally observed, predicted, tissue-specific, or disease-associated isoforms. Primer performance may also depend on polymerase, buffer composition, ionic conditions, annealing temperature, template abundance, and local sequence context. Accordingly, the approximately 15 °C criterion should be viewed as an experimentally supported design rule rather than an absolute boundary under every PCR condition, and empirical validation remains important. In cases not resolved by junction-specific PCR, complementary approaches such as long-read sequencing, length-dependent PCR, or RACE may be required.

An additional advantage of exon–exon junction (EEJ)-guided primer design is its ability to reduce heteroduplex formation during simultaneous amplification of highly similar transcript variants. In the HTRA1-AS1 model, Sanger sequencing and denaturation–reannealing experiments confirmed that unexpected intermediate products were heteroduplexes formed between ENST416 and HTRA1-AS1 amplicons. Additional higher-molecular-weight bands were also observed, consistent with the formation of higher-order heteromeric complexes through interactions between partially homologous PCR products. Although G-quadruplex and other higher-order DNA structures have been reported, to our knowledge, higher-order complexes generated by intermolecular assembly of partially homologous PCR amplicons have not been previously described [32, 33]. The exact molecular architecture of these higher-order products, however, requires further structural characterization.

Importantly, heteroduplex formation was reduced by using variant-specific primers, including an EEJ primer, to decrease homologous sequence overlap between amplicons. Thus, EEJ-guided primer design offers an advantage beyond transcript specificity by reducing intermolecular hybridization and improving the interpretability of transcript-variant RT-PCR. In summary, experimentally defined Tm-guided EEJ primer design provides a practical strategy for distinguishing highly similar RNA transcript variants when sufficiently large transcript-specific exonic regions are unavailable. By constraining the Tm of each exon-matching arm relative to the PCR annealing temperature while preserving stable full-length binding across the intended junction, the approach favors junction-dependent amplification and reduces amplification arising from partial primer binding. The experimentally supported arm-specific Tm criterion therefore provides a quantitative framework for designing transcript-specific RT-PCR and qPCR assays based on exon connectivity.

## Data Availability

The genome-wide transcript characterization reported in this study was performed using a fixed RefSeq dataset retrieved on June 1, 2026. Processed data underlying the transcript characterization and primer-design analyses are provided in the Supplementary Data.

## Author contributions

Initiative and conceptualization: PWZ Methodology and analysis design: JN, YL, PWZ Data collection and analysis: JN, YL, PWZ, DJZ Writing: JN, YL, PWZ, DJZ

