## supplementary file for "Experimentally Defined Tm-Guided Exon–Exon Junction Primer Design Enables Specific Discrimination of Highly Similar RNA Variants"

**Supplementary Data**

1. Variant-specific PCR Primer Information

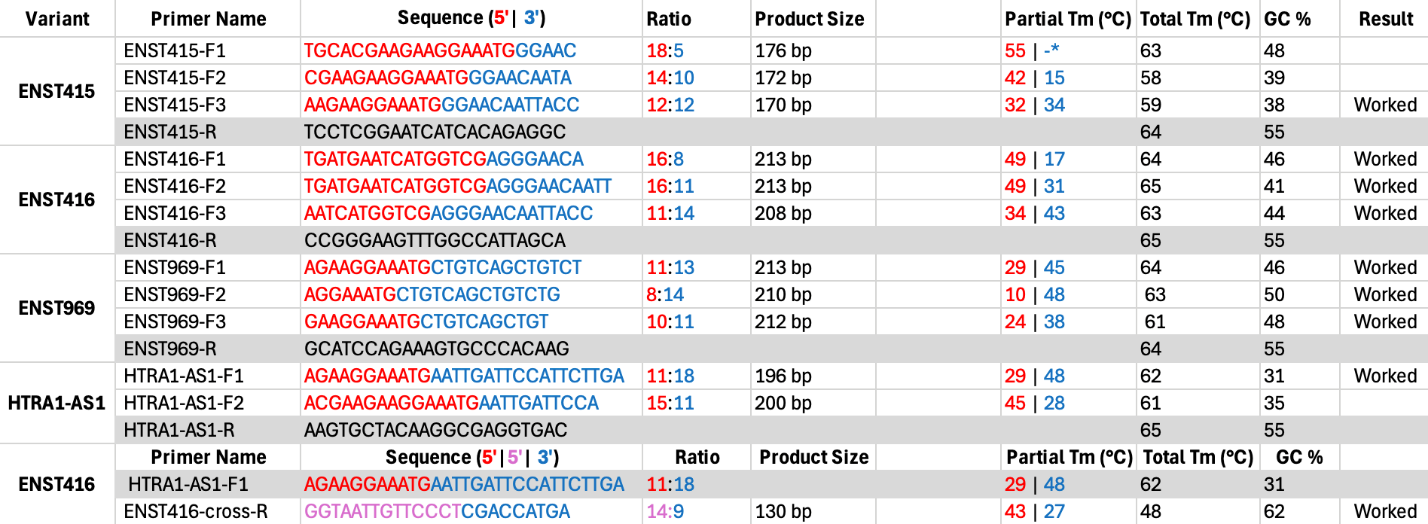

2. Variants Combined PCR Primer Information

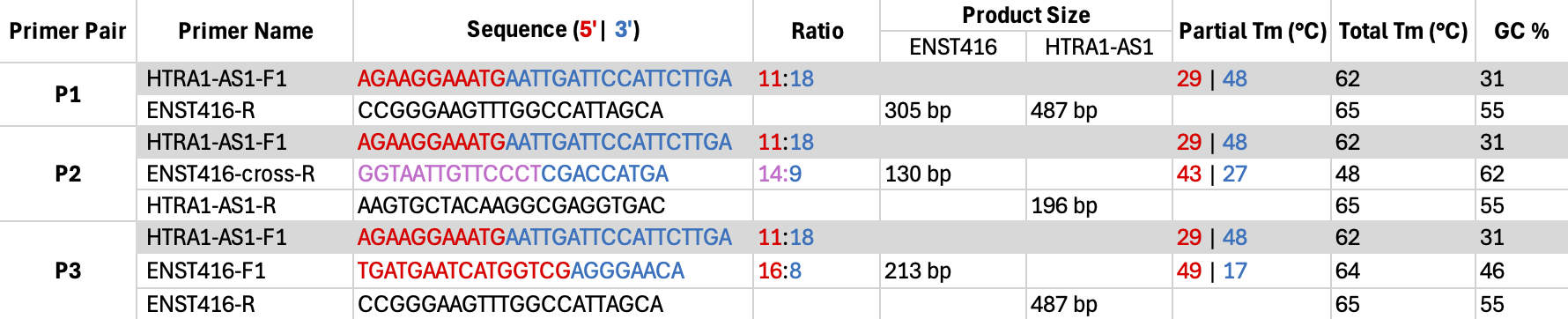

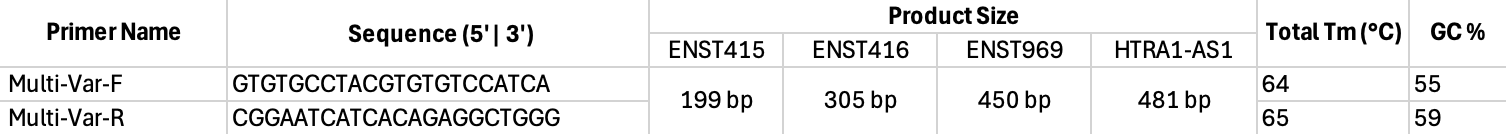

3. PCR Protocol and Reagent Information

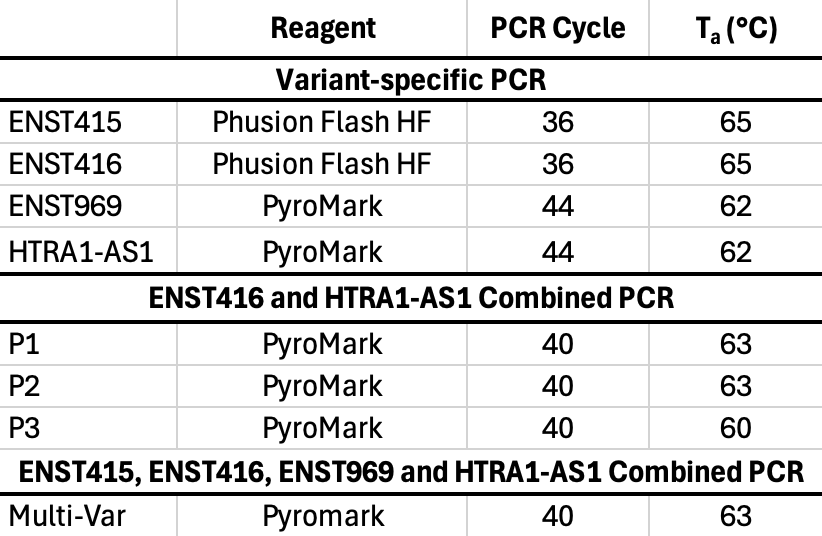

4. Gel Electrophoresis Image and Information of transcript-specific PCR products from GAPDH and APEX1

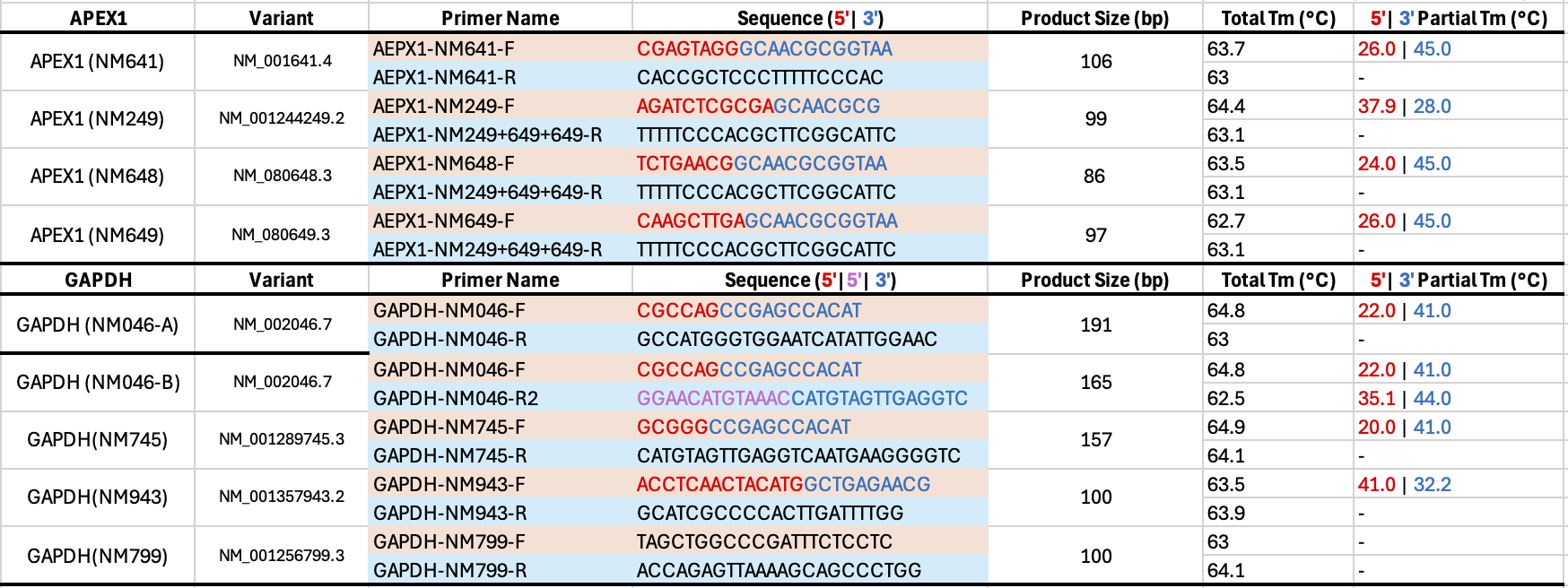

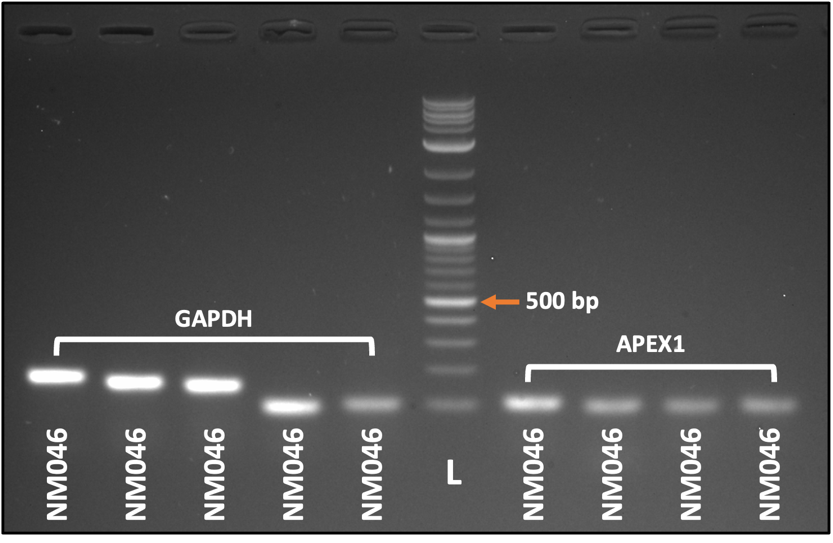

5. Sanger Sequencing Data of the GAPDH and APEX1

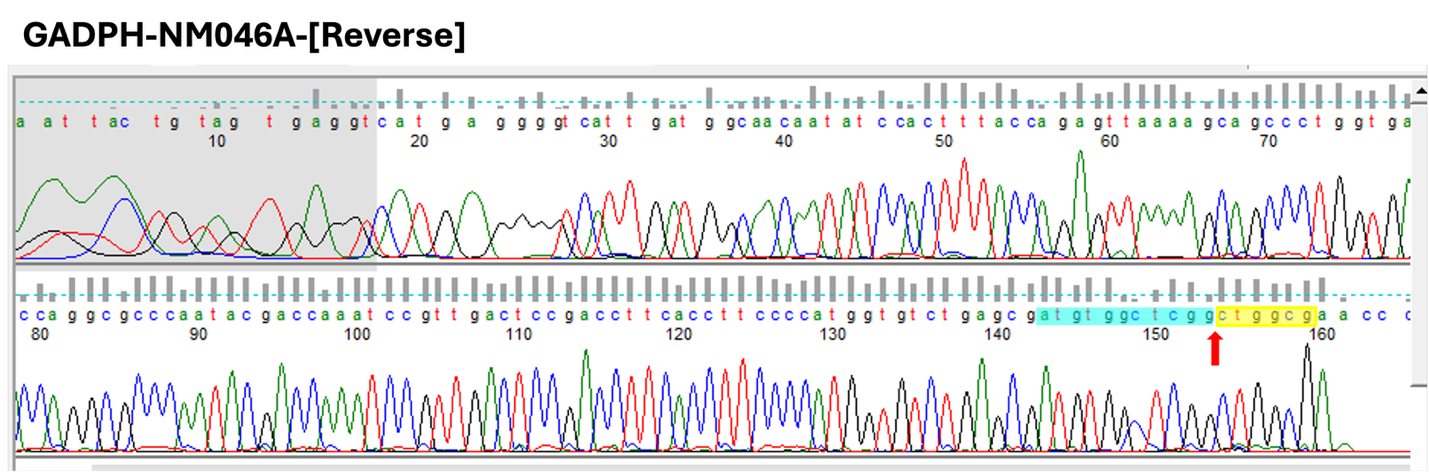

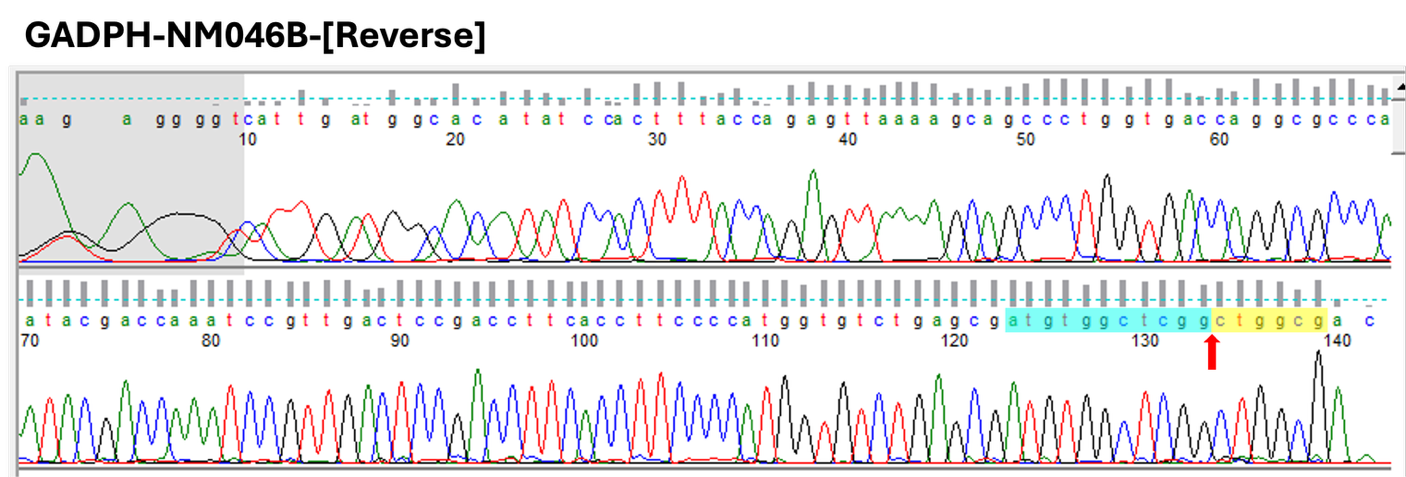

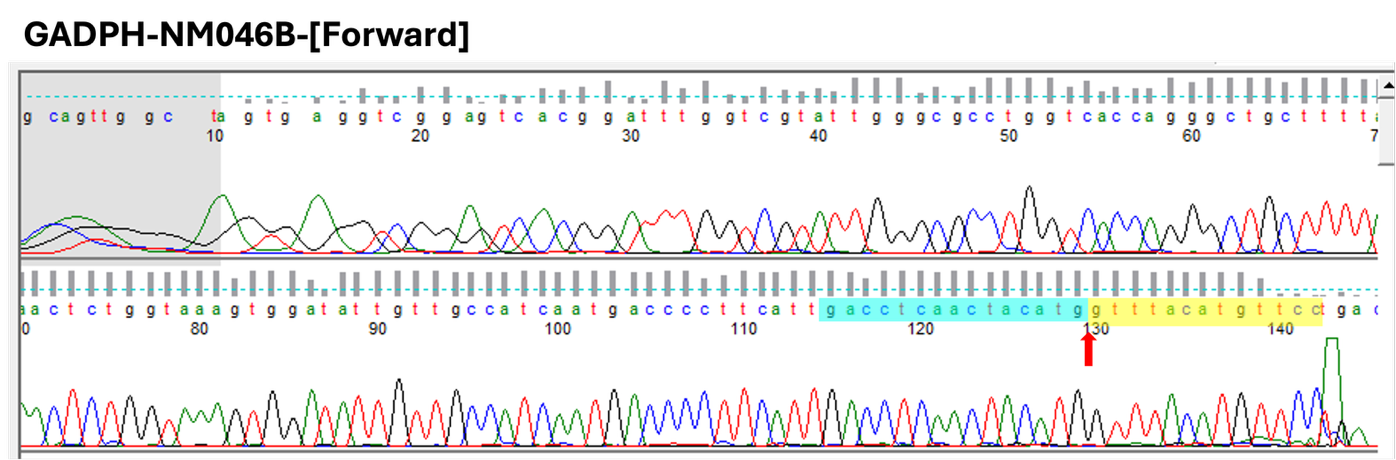

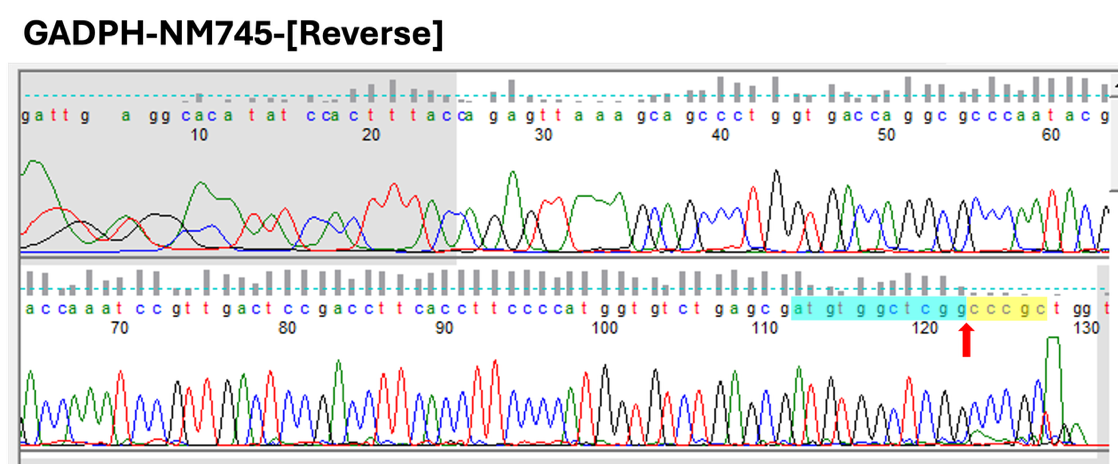

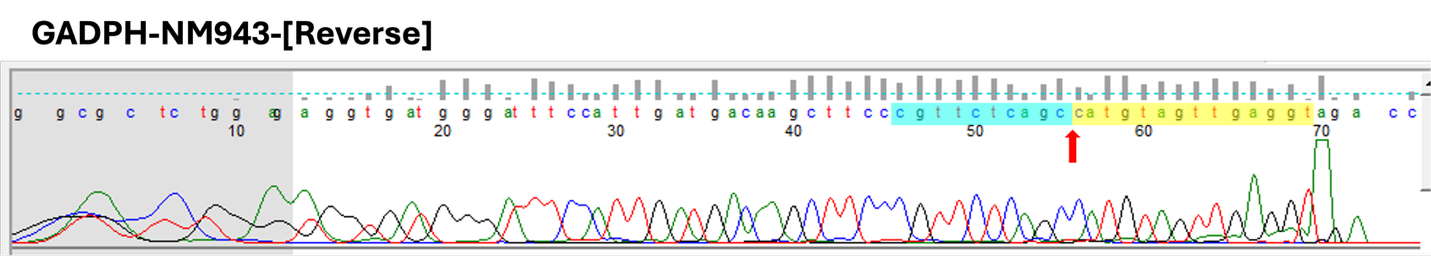

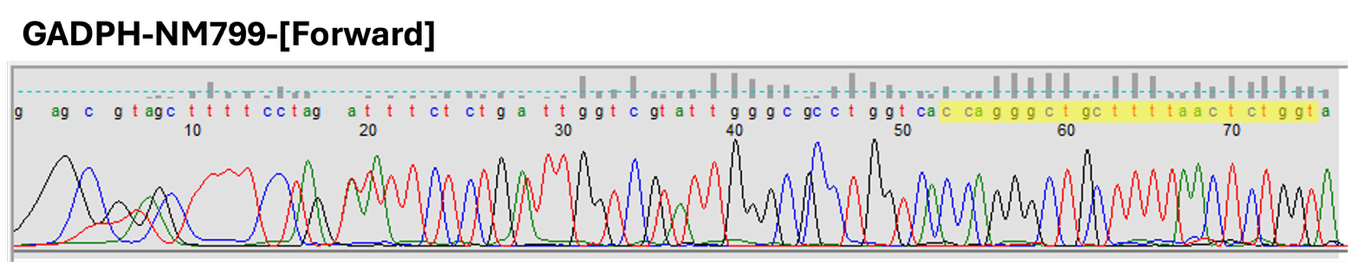

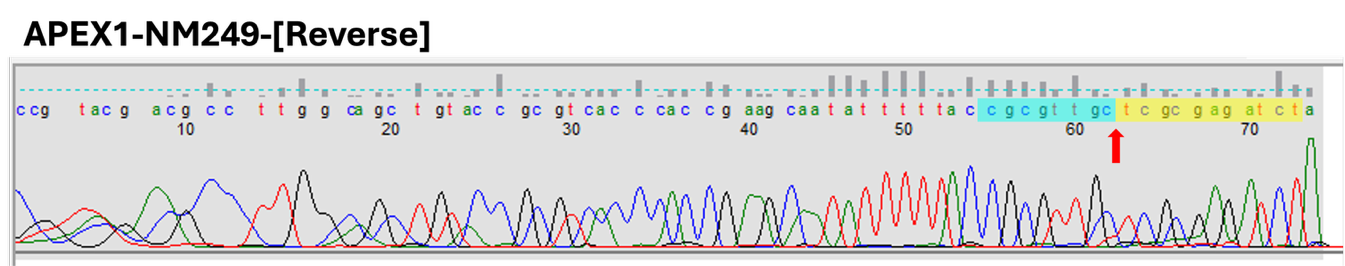

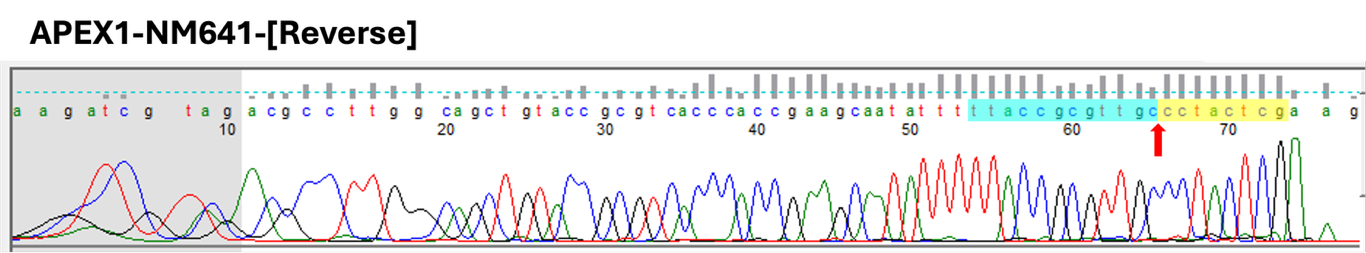

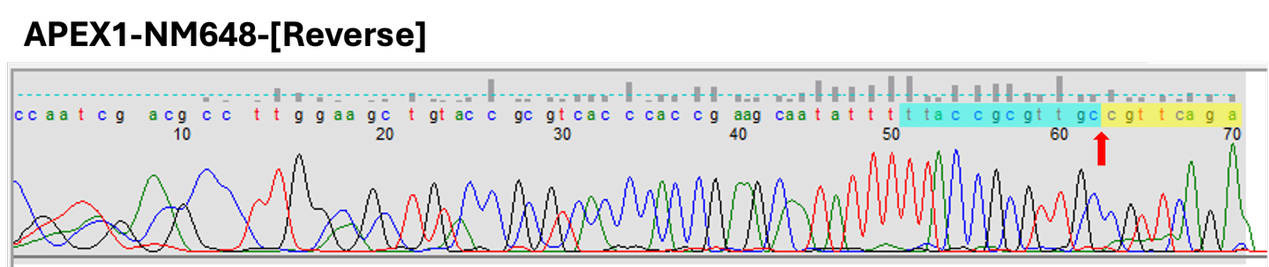

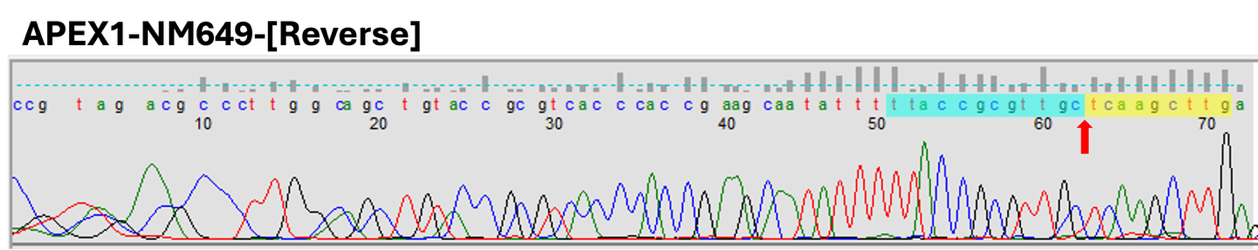

6. Performance Comparison of the Primer Design Tools

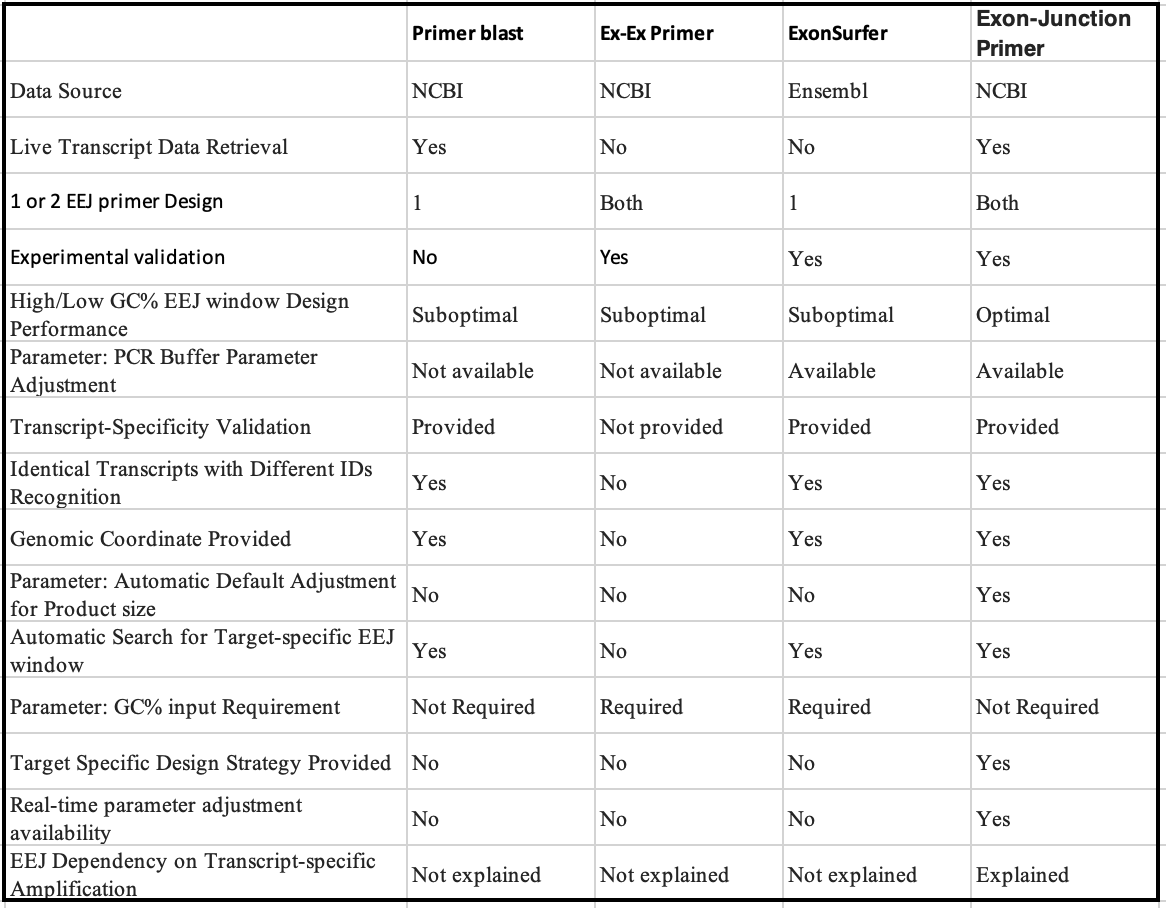
